# hL1 retrotransposons drive human neuronal transcriptome complexity and functional diversification

**DOI:** 10.1101/2023.03.04.531072

**Authors:** Raquel Garza, Diahann Atacho, Anita Adami, Patricia Gerdes, Meghna Vinod, PingHsun Hsieh, Ofelia Karlsson, Vivien Horvath, Pia A. Johansson, Ninoslav Pandiloski, Jon Matas, Annelies Quaegebeur, Antonina Kouli, Yogita Sharma, Marie E Jönsson, Emanuela Monni, Elisabet Englund, Evan E. Eichler, Molly Hammell, Roger A. Barker, Zaal Kokaia, Christopher H. Douse, Johan Jakobsson

## Abstract

The genetic mechanisms underlying the expansion in size and complexity of the human brain remains poorly understood. L1 retrotransposons are a source of divergent genetic information in hominoid genomes, but their importance in physiological functions and their contribution to human brain evolution is largely unknown. Using multi-omic profiling we here demonstrate that L1-promoters are dynamically active in the developing and adult human brain. L1s generate hundreds of developmentally regulated and cell-type specific transcripts, many which are co-opted as chimeric transcripts or regulatory RNAs. One L1-derived lncRNA, LINC01876, is a human-specific transcript expressed exclusively during brain development. CRISPRi-silencing of LINC01876 results in reduced size of cerebral organoids and premature differentiation of neural progenitors, implicating L1s in human-specific developmental processes. In summary, our results demonstrate that L1-derived transcripts provide a previously undescribed layer of primate- and human-specific transcriptome complexity that contributes to the functional diversification of the human brain.

## Introduction

During evolution, primate brains have expanded in size and complexity resulting in a unique level of cognitive functions. The genetic alterations responsible for this enhancement remain poorly understood ^2–5^. Our closest living relative, the chimpanzee, shares more than 98% of protein-coding sequences with humans – making it unlikely that species-specific protein-coding variants are the sole evolutionary drivers of brain complexity ^6, 7^. Rather, a significant fraction of the genetic basis for the differences in non-human primate and human brains likely reside in the non-coding part of the genome.

A large portion of genetic information specific to primates is stored in transposable elements (TEs), mobile genetic elements that make up almost 50% of the human genome^1^. Since TEs have populated the genome through mobilization this has resulted in major inter-species and inter-individual differences in their genomic composition. Hundreds of thousands of TEs are primate-specific and several thousand of them are human-specific ^8, 9^. TEs pose a threat to genomic integrity – as their activation may result in retrotransposition events that cause deleterious mutations ^10, 11^ – and the host has therefore evolved numerous mechanisms to prevent mobilization ^12, 13^. In somatic human tissues such as the brain, it is thought that the vast majority of TEs are transcriptionally repressed, which correlates with the presence of DNA CpG-methylation ^14, 15^. However, TEs have the potential to be exapted – providing a benefit for the host as a source of gene regulatory elements and co-opted RNAs and peptides ^16^. For example, TEs are largely responsible for the emergence of species-specific long non-coding RNAs (lncRNA).^17^, which are non-translated transcripts of more than 200 nucleotides that have been implicated to control a wide variety of cellular processes ^18^.

The most abundant and only autonomously-mobilizing TE family in humans is long interspersed nuclear element-1 (L1) ^19^. The human genome holds around half a million individual L1 copies, occupying ∼17% of genomic DNA, including ancient fragments and evolutionarily younger full-length copies ^1, 20^. Since L1s have colonized the human genome via a copy-and-paste mechanism in different waves, it is possible to approximate the evolutionary age of each individual L1 copy and assign them to chronologically-ordered subfamilies ^21^. Only full-length L1s (>6kbp) with an intact 5′ UTR allows for element-derived expression. However, most L1s are inactivated due to 5’ truncations and the accumulation of inactivating deletions and mutations. Full-length L1s are transcribed from an internal 5’ RNA polymerase II promoter as a bicistronic mRNA encoding two proteins, ORF1p and ORF2p, which are essential for L1 mobilization ^22–24^. Notably, the L1 promoter is bidirectional and in evolutionarily-young L1s the antisense transcript encodes a small peptide, ORF0, with poorly characterized function ^25, 26^. L1-antisense transcripts can also give rise to chimeric transcripts and act as alternative promoters for protein coding genes ^14, 26^.

Over the last two decades L1 activity has been implicated in the functional regulation of the human brain, primarily based on the observation of somatic L1 retrotransposition events in the neural lineage leading to genomic mosaicism ^27–33^. However, it has been challenging to determine the functional impact of such events, which are rare and randomly distributed. Given their abundance and repetitive nature, L1s are difficult to study using standard molecular biology techniques. For example, estimation of L1-derived RNA expression using quantitative PCR-based techniques or standard short-read RNA sequencing (RNA-seq) approaches, whether bulk or single-cell, often fail to separate L1 expression originating from the L1 promoter from that of bystander transcripts that are the result of readthrough transcription ^34^. Therefore, it is still debated if and in which cell types L1 expression occurs in the developing and adult human brain and the impact of L1s on the physiology of the human brain remains unresolved.

In this study we have used a combination of bulk short-read, long-read and single-nuclei RNA-seq coupled with CUT&RUN epigenomic profiling, together with tailored bioinformatical approaches ^35, 36^ to demonstrate that L1-derived transcripts are highly expressed in the healthy developing and adult human brain. We found that the bidirectional L1 promoter is dynamically active, resulting in the generation of hundreds of L1-derived transcripts that display developmental regulation and cell-type specificity. We provide evidence for the expression of full-length L1s as well as L1s that are co-opted as regulatory RNAs or alternative promoters. One human-specific L1-derived lncRNA (L1-lncRNA), LINC01876, is exclusively expressed during human brain development. CRISPRi-based silencing of LINC01876 results in reduced size of cerebral organoids and premature differentiation of neural progenitor cells (NPCs) and neurons – suggesting that it has an important role in brain development. Together, these results demonstrate that L1-derived transcripts are abundant in the human brain where they provide an additional layer of primate- and human-specific transcriptome complexity that has contributed to the evolution of the human brain.

## Results

### L1-derived transcripts are abundant in the adult human brain

To investigate the expression of L1s in the adult human brain we obtained cortical tissue biopsies (temporal and frontal lobe) from three non-neurological deaths in people aged 69, 75 and 87 years. We sorted cell nuclei from the biopsies, extracted RNA and used an *in-house* 2x150bp, polyA-enriched stranded library preparation for bulk RNA-seq using a reduced fragmentation step to optimize library insert size for L1 analysis. Such reads can be mapped uniquely and assigned to individual L1 loci, except for reads originating from a few of the youngest L1s and polymorphic L1 alleles that are not in the hg38 reference genome. We obtained ∼30 million reads per sample. To quantify L1 expression we used two different bioinformatic methodologies (Figure 1A). First, we allowed reads to map to different locations (multi-mapping) and used the *TEtranscripts* software ^35^ in multi-mode to best assign these reads (Supplemental Figure 1A). Second, we discarded all ambiguously mapping reads and only quantified those that map uniquely to a single location (unique mapping).

**Figure 1.**
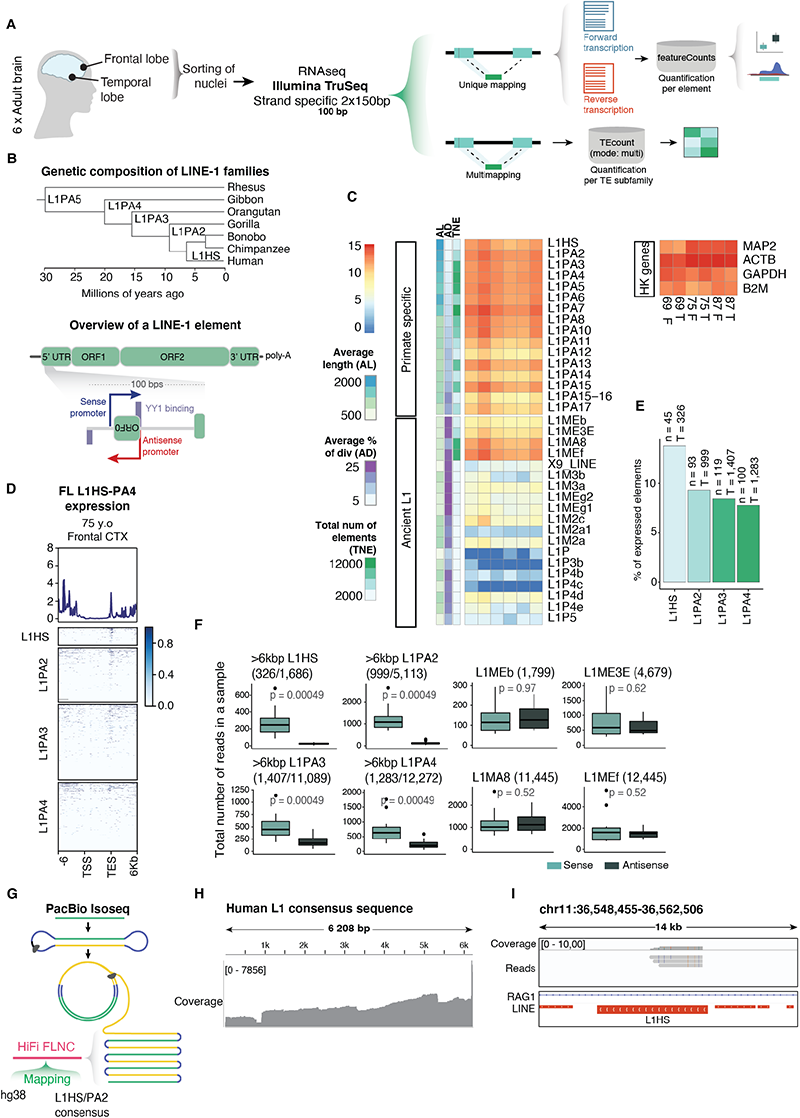
L1-derived transcripts are abundant in the adult human brain. A) Schematic illustrating sample collection, sequencing strategy and bioinformatical approach. B) Top: Phylogenetic tree showing the evolutionary age of young L1 subfamilies. Bottom: Structure of a L1 element with a zoom-in to its 5’ UTR. Arrows indicate promoters in sense (blue) and antisense (red). YY1 binding sites indicated in purple boxes (sense on top, antisense on bottom). C) Expression of primate-specific L1 subfamilies compared to ancient L1 subfamilies and selected housekeeping genes as reference. Row annotation showing average length (AL), average percentage of divergence from consensus (AD), and the total number of elements (TNE) (information extracted from RepeatMasker open-4.0.5). D) Expression (RPKM) over full-length (>6kbp) L1HS, L1PA2, L1PA3 and L1PA4, plus 6kbp flanking regions. E) Percentage of expressed full-length (>6kbp) elements (mean normalized counts > 10; see methods) among young L1 subfamilies (n = number of expressed elements; T= total number of full-length elements). F) Read counts in sense (light teal) and antisense (dark teal) per sample. First four showing full-length elements in young L1 subfamilies, last four showing ancient L1 subfamilies with a comparable number of copies. G) PacBio Iso-Seq schematic and mapping approach. H) Coverage of PacBio Iso-Seq library mapped to L1HS and L1PA2 consensus sequence. I) Genome browser tracks showing PacBio Iso-Seq reads over the promoter region of a full-length L1HS.

We found that L1s expressed in the adult human brain primarily belonged to primate-specific families, including both hominoid-specific (L1PA2 – L1PA4) and human-specific elements (L1HS) (Figure 1B) ^21^. The total expression level of these subfamilies, as quantified with *TEtranscripts* ^35^, corresponds to expression levels of housekeeping genes (Figure 1C). Using unique mapping, we were able to detect expression coming from hundreds of evolutionarily young L1s (Figure 1D), including 138 full-length L1HS or L1PA2 elements (Figure 1E). The RNA-seq signal over the full-length L1s was highly enriched at the 3’-end, which reflects the presence of degraded RNA in human post-mortem samples and L1-mappability issues in the central part of the element, but also indicates that the transcription of L1s terminate in the internal L1 polyadenylation signal ^37^. Importantly, when comparing the number of reads transcribed in the same orientation as the L1s (in sense) to those in the opposite direction (in antisense), we found that most of the transcription in these regions was in sense to the L1s (Figure 1F and Supplemental Figure 1B). This suggests that most L1-transcripts originates from the L1-promoter and are not a consequence of read-through or bystander transcription. In a few cases, we also found clear evidence of activity of the antisense L1-promoter ^26^, resulting in transcription extending out into the upstream flanking genome (Supplemental Figure 1C).

To complement this analysis, we performed long-read PacBio Iso-Seq on a cortical biopsy from a deceased 63-year-old man (Figure 1G). This allows for the identification of L1-derived transcripts that can be accurately mapped to full-length L1s and enables the identification of TSSs and splicing events. We mapped reads (mean read length 2kbp) to the L1HS and L1PA2 consensus sequence to which 10.9K reads mapped (of a total of 2.9M reads in the library). The density of the mapped reads throughout the sequence reflected the common 5’ truncation that is present in most L1 copies in the human genome ^20, 38^, but 1,360 reads still mapped to the 5’UTR (Figure 1H). Notably, we found several clear examples of long reads mapping to the promoter region of young full-length L1s providing further support to L1 promoter-driven expression in the adult human brain (Figure 1I).

### L1 expression is enriched in neurons in the adult human brain

To investigate the expression of L1s at cell-type resolution, we performed snRNA-seq analysis using the 3’ 10X Chromium Platform and five of the adult cortical samples we sequenced in bulk RNAseq (Figure 2A). In total, we sequenced 8,089 high-quality nuclei with a mean of 3,042 genes detected per cell. Unbiased clustering using Seurat resulted in 22 clusters (Figure 2B) and based on the expression of canonical gene markers we identified excitatory neurons, inhibitory neurons, astrocytes, oligodendrocytes, oligodendrocyte precursors and microglia at expected ratios (Figure 2C-D and Supplemental Figure 2A).

**Figure 2.**
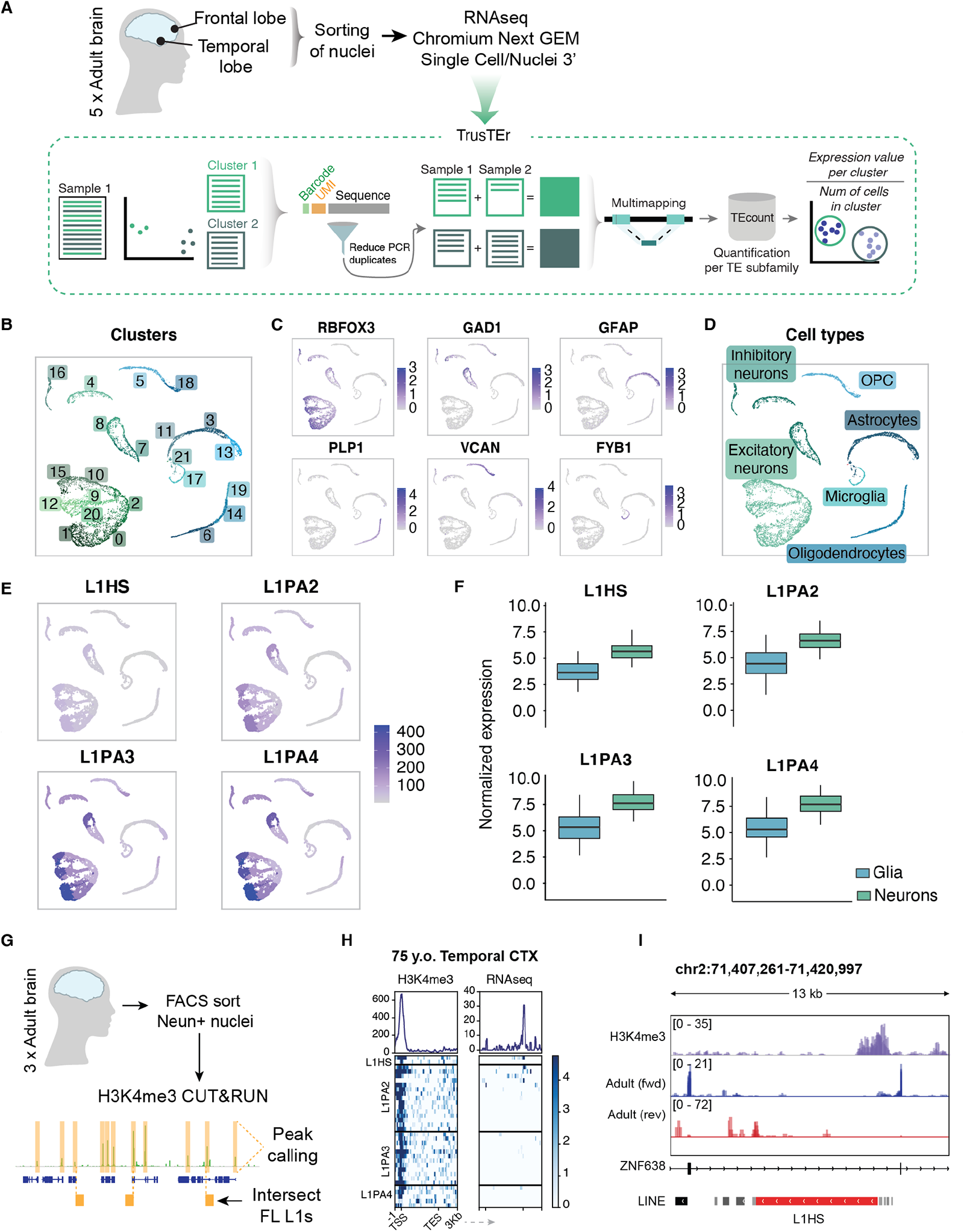
L1 expression in neurons in the adult human brain. A) Schematic of sample collection, sequencing approach and analytical bioinformatics pipeline for TE expression in single-nuclei data. B) Single nuclei RNAseq: UMAP coloured by defined clusters. C) Expression of selected markers for different cell types. D) UMAP coloured by characterized cell types. E) Pseudo-bulk cluster expression of young L1 subfamilies on UMAP. F) Comparison of glia vs neuronal clusters per L1 family. G) Schematic of NeuN+ H3K4me3 CUT&RUN in adult human brain samples and bioinformatical approach. H) H3K4me3 peaks (left heatmap) over full-length L1 subfamilies (L1HS – L1PA4) and RNAseq signal (right heatmap). Profile plots showing sum of signal. I) Genome browser tracks showing the expression of a full-length L1HS with an H3K4me3 peak on its promoter and RNAseq signal (RPKM) split by direction of transcription (blue = forward; red = reverse).

Quantification of L1 expression is challenging using single-cell technologies, as the number of mapped reads in a single cell falls short of accurate quantification, regardless of the mapping technique. To circumvent this limitation, we used an *in-house* bioinformatic pipeline allowing the analysis of L1 expression from the snRNA-seq dataset (Figure 2A). This method uses the cell clusters determined based on gene expression. Then, by back-tracing the reads from cells forming each cluster, it is possible to analyze the expression of L1s, using the *TEtranscripts* software ^39^ or with unique-mapping, in distinct cell populations. This pseudo-bulk approach greatly increases the sensitivity of the TE analysis and enables quantitative estimation of L1 expression at single-cell-type resolution ^36^.

We found clear evidence of L1 expression in the snRNA-seq data. Notably, L1 expression was higher in neurons, including both excitatory and inhibitory neurons, when compared to different glial cell types (Figure 2E-F and Supplemental Figure 2B). To confirm that L1s were expressed in neurons, but not in glia, we examined the transcription of each cluster per individual element using unique mapping (Supplemental Figure 2B). Profile plots on reads from neurons displayed distinctive peaks over the elements, which correlated with the mappability of the L1s (Supplemental Figure 2B-D). In line with the bulk RNA-seq data, we observed that L1 expression was confined to evolutionary young elements and that the antisense signal over L1HS and L1PA2s was negligible, implying that the signal in sense of the elements is not due to readthrough or bystander transcription (Supplemental Figure 2C).

To further confirm that the L1 expression in human neurons originates from the L1 promoter we performed 5’-enriched snRNA-seq using the 10X Chromium Platform since this approach allows detection of the TSSs (Supplemental Figure 2E). We again observed expression of evolutionary young L1s in neurons but not in glia (Supplemental Fig 2G-H) further strengthening the observation that L1-expression in human neurons originates from the L1 promoter.

### Identification of H3K4me3 at the L1 promoter in adult human neurons

The bulk and snRNA-seq analyses demonstrate that L1s are highly expressed in adult human neurons. However, due to the presence of many polymorphic L1s in the human population it is not possible to assign this expression to individual elements with complete certainty due to the absence of such polymorphic L1s in the hg38 reference genome ^9, 40^. To address this issue, we performed CUT&RUN epigenomic analysis ^41^ on adult human neurons to identify if the histone mark H3K4me3, which is associated with active promoters, was present on L1s. Since the signal of this histone modification spreads to the unique flanking genomic context, this approach allows for an accurate identification of transcriptionally-active individual L1 loci ^14^. To this end, we FACS-isolated neuronal nuclei (NeuN+) from the same three human cortical biopsies used for the transcriptomic analysis and performed CUT&RUN. The resulting sequencing data were uniquely mapped, followed by peak calling and intersection with full-length L1s (Figure 2G).

The H3K4me3 analysis identified 38 high-confidence H3K4me3 peaks located in the TSS of full-length evolutionary young L1s (Figure 2H) (several elements were confirmed to be expressed in the bulk RNA-seq dataset). These elements represent examples of L1 transcriptional activity in adult human neurons that can be *bona fide* assigned to individual elements. For example, we found a full-length L1HS located in the intron of ZNF638 as being transcriptionally active in adult human neurons (Figure 2I).

### L1s are expressed during human brain development

To investigate whether L1s are also expressed during human brain development, we analyzed six human fetal forebrain samples aged 7.5 – 10.5 weeks post-conception using our multi-omics approach (Figure 3A, Supplemental Fig 3A). The bulk RNA-seq analysis demonstrated that evolutionary young L1s are expressed at levels approaching those of housekeeping genes in forebrain human development (Figure 3B). We found no obvious difference in the magnitude of expression between the different gestational ages of the tissue. Unique mapping revealed that hundreds of different L1 loci were expressed, with the majority of these displaying sense strand enrichment, indicating an active L1 promoter (Figure 3C-E, Supplemental Fig 3B-C). In line with this, the H3K4me3 analysis confirmed that several L1s carried this histone mark over the TSS, thus representing *bona fide* examples of unique L1 integrant expression in brain development (Fig 3F). We also found evidence of antisense transcription initiated at the L1 TSS to the upstream genome (Supplemental Figure 3D).

**Figure 3.**
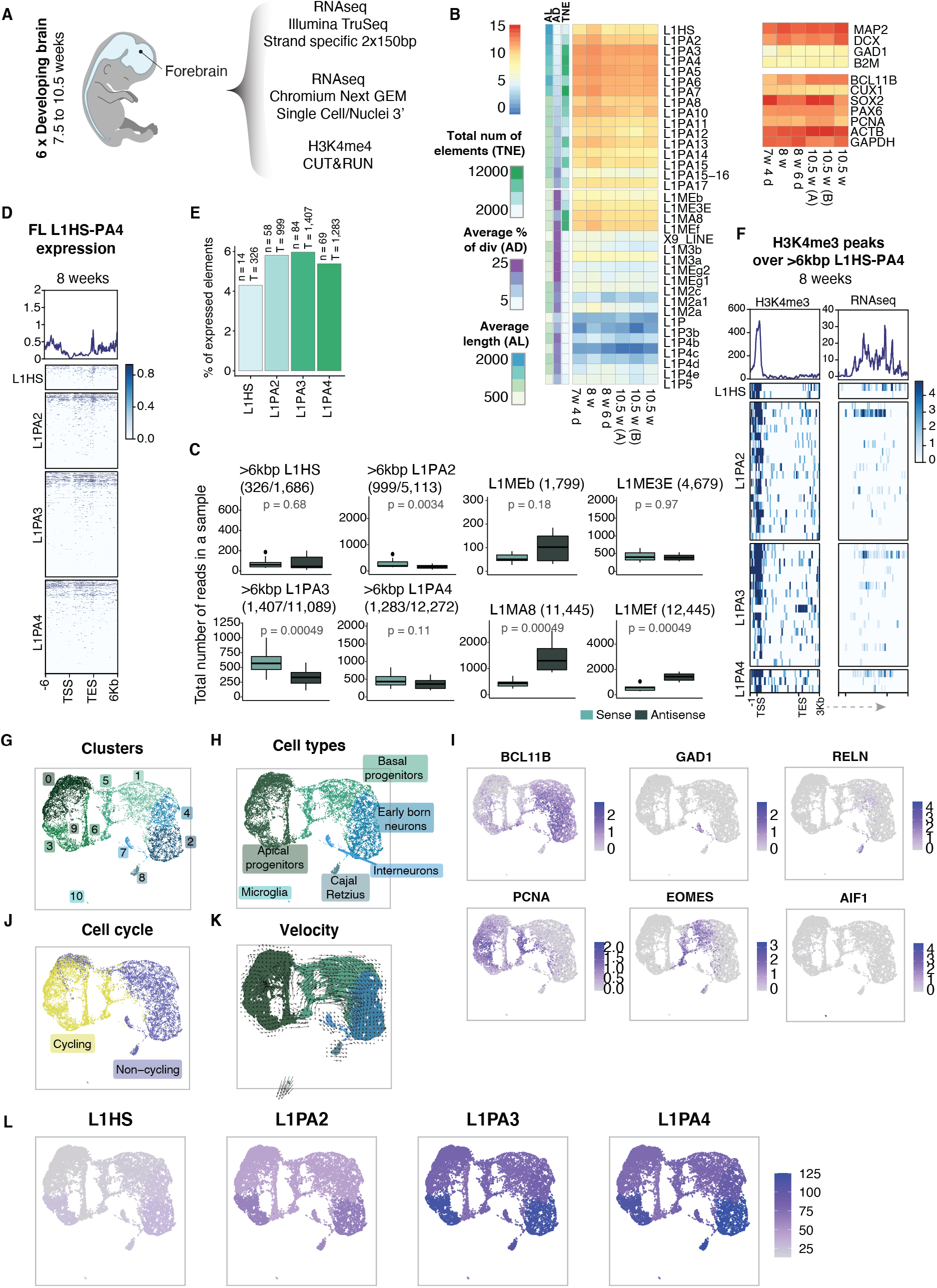
L1s are expressed in human brain development. A) Schematic of sequencing strategy of fetal human forebrain samples B) Expression of primate specific L1 subfamilies compared to ancient L1 subfamilies, and selected housekeeping and development-related genes as reference. Row annotation showing average length (AL), average percentage of divergence from consensus (AD), and the total number of elements (TNE) (information extracted from RepeatMasker open-4.0.5). C) Read count in sense (light teal) and antisense (dark teal) per sample. First four boxplots showing full-length elements in young L1 subfamilies, last four showing ancient L1 subfamilies with a comparable number of copies. D) Expression (RPKM) over full length (>6kbp) L1HS, L1PA2, L1PA3 and L1PA4, plus 6kbp flanking regions. E) Percentage of expressed full length (>6kbp) elements (mean normalized counts > 10; see methods) among young L1 subfamilies (n = number of expressed elements; T= total number of full-length elements). F) Detected H3K4me3 peaks (left heatmap) over full length L1 subfamilies (L1HS till L1PA4) and RNAseq signal (right heatmap). Profile plots showing sum of signal. G) Fetal human forebrain single nuclei RNAseq UMAP colored by cluster. H) UMAP colored by cell types. I) Expression of selected biomarkers for different cell types. J) UMAP colored by cell cycle state (based on CellCycleScoring from Seurat). K) Velocity plot colored by cell type. L) Pseudo-bulk cluster expression of young L1 subfamilies on UMAP.

A notable difference when comparing the data from development to the adult brain was the expression of L1HS, which are human-specific L1s of which some retain the capacity to retrotranspose ^19, 42^. When analyzing strand-specific expression in the developing brain samples we found no enrichment for sense strand expression of L1HS and we found very few L1HS expressed among the elements detected from the different subfamilies (Figure 4C&E). This contrasts with the adult samples where we detected clear evidence for sense-strand expression of L1HS expression and found many unique L1HS loci to be expressed (Figure 1E). Thus, L1HS expression, which includes all elements with retrotransposition capacity ^19, 42^, appears to be selectively silenced during human brain development.

**Figure 4.**
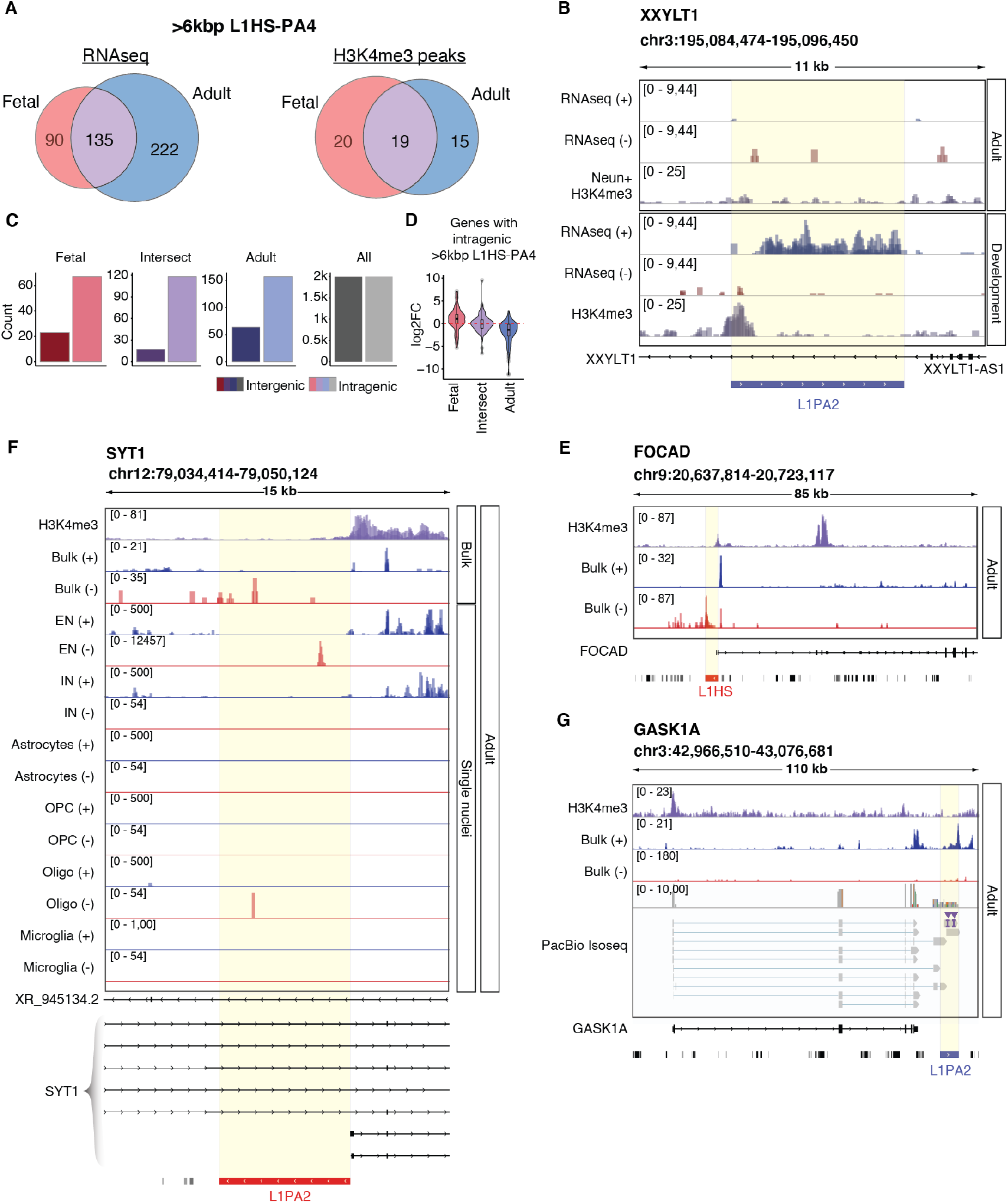
L1s are dynamically expressed in the developing and adult human brain. A) Left: Number of expressed L1HS-L1PA4 (>6kbp) in fetal (red) and adult samples (blue) (mean normalized counts > 10; see methods), and the number of elements found to be expressed in both datasets (intersection; purple). Right: Number of H3K4me3 peaks over L1HS-L1PA4 (>6kbp) in fetal (red) and adult samples (blue) and the intersection between datasets (purple). B) Genome browser track showing the expression of a development-specific full-length L1PA2 with an H3K4me3 peak at its promoter. C) Histograms showing the number of intragenic (light) or intergenic (dark) L1HS-L1PA4s (>6kbp) in fetal (red), adult (blue) or those expressed in both datasets (purple). D) log2FoldChange of the genes with an intragenic L1HS-L1PA4 (>6kbp) in fetal (red), adult (blue) and the intersection (purple) (fetal vs adult (ref); DESeq2). Genome browser tracks showing (from top to bottom): NeuN+ H3K4me3 CUT&RUN of adult samples (samples overlayed in purple), bulk RNAseq of adult samples (overlayed) split by strand (blue = forward; red = reverse) over E) FOCAD’s transcription starting site with an antisense full length L1HS. F) SYT1 with an antisense full length L1PA2 at the beginning of one of its isoforms. Additional tracks showing overlayed cluster expression (adult single nuclei RNAseq) of excitatory neurons (EN), inhibitory neurons (IN), Astrocytes, Oligodendrocyte precursor cells (OPCs), Oligodendrocytes (Oligo), and Microglia. G) GASK1A with an L1PA2 downstream of the gene’s transcription ending site. Bottom panel shows PacBio Iso-Seq, validating the formation of a new GASK1A isoform with the L1PA2 incorporated.

We performed snRNA-seq on the fetal forebrain samples and sequenced 12,183 high-quality nuclei with a mean of 3,818 genes detected per cell. Unbiased clustering using Seurat resulted in 11 clusters (Figure 3G) and based on the expression of canonical gene markers representing cell types present at this developmental stage we identified apical progenitors, basal progenitors, early-born neurons, immature interneurons, Cajal Retzius cells, and microglia (Figure 3H-I). We also used RNA velocity ^43^ and scoring of cell-cycle related genes to further characterize this dataset (Figure 3J-K). These analyses revealed, in line with the existing literature, that apical progenitors represent an early proliferative neural progenitor stage that with time is replaced by more mature basal progenitors and post-mitotic immature neurons ^44^. L1 expression levels were similar in apical progenitors, basal progenitors and early-born neurons (Figure 3L and Supplemental Figure 3E-F) and we found no significant correlation between L1 expression level and cell-cycle state (Supplemental Figure 3G). Thus, L1s are expressed in human forebrain development already at the progenitor stage and expression is not substantially increased upon differentiation and exit of the cell cycle.

### Individual L1 loci are dynamically expressed in the developing and adult human brain

Our results demonstrate that the internal L1 promoter is active in the developing and adult human brain resulting in the transcription of a wide panel of L1-derived transcripts. However, we noted that the developing and adult brain samples distinctly differed in the expression of individual L1 loci. When we intersected RNA-seq or H3K4me3 data from the developing and adult brain we found that only a minority of L1 loci were expressed in both sample types (Figure 4A). For example, we found a full-length L1PA2 on chromosome 3 that was highly expressed in brain development but completely silent in the adult brain (Figure 4B). Thus, the bulk of the L1 expression from unique elements was either confined to development or the adult brain indicating that the expression of different L1 loci depends on cellular context ^45^.

Since individual L1 loci share the same regulatory sequences, we hypothesized that the divergent expression in the developing and adult brain is a consequence of the L1-integration site and the transcriptional activity of the nearby genome. In line with this, we noted that expressed L1 loci were highly enriched to intragenic regions (Figure 4C). Notably, the expression of the genes in these regions clearly correlated with the expression of individual L1 loci (Figure 4D). For example, L1s expressed uniquely during development were often located in introns of genes with a developmental specific expression pattern (Figure 4D). Thus, this analysis indicates that expression of individual L1 loci is governed by their integration site and the transcriptional activity of the nearby genome.

### L1-derived transcripts contribute to transcriptome complexity in human neurons

The activity of the L1 promoter in the human brain suggests that L1s are a rich potential source of primate-specific and human-specific transcripts, that in turn may be co-opted and contribute to transcriptome complexity and speciation. When searching our dataset, we found several such examples of co-option where L1s appear to have integrated into and modified the human transcriptome. For example, an L1HS in the *FOCAD* gene acts as an alternative promoter of this gene (Figure 4E). Likewise, an L1PA2 provides an alternative promoter for an isoform of SYT1, which is exclusively expressed in neurons (Figure 4F). The long-read RNA-seq analysis in combination with bulk RNA-seq identified an L1PA2 acting as an alternative 3’ UTR in GASK1A (Figure 4G). Thus, our multi-omics approach revealed several novel examples where L1s are integrated into the gene regulatory landscape of the developing and mature human brain. Notably, all these L1s represent hominoid- or human-specific insertions.

To investigate the potential role of L1s in contributing to normal human brain functions we focused on a transcriptionally active full-length L1PA2 element on chromosome 2 (6013 bp long). The L1 antisense promoter ^14, 26^ serves as the TSS of a lncRNA: LINC01876. RNA-seq, snRNA-seq and H3K4me3-CUT&RUN supported that the L1PA2 act as an antisense promoter for this L1-lncRNA in human brain development (Figure 5A). Notably, this expression appears to be limited to development since no LINC01876 expression was found in the adult brain (Figure 5A).

**Figure 5.**
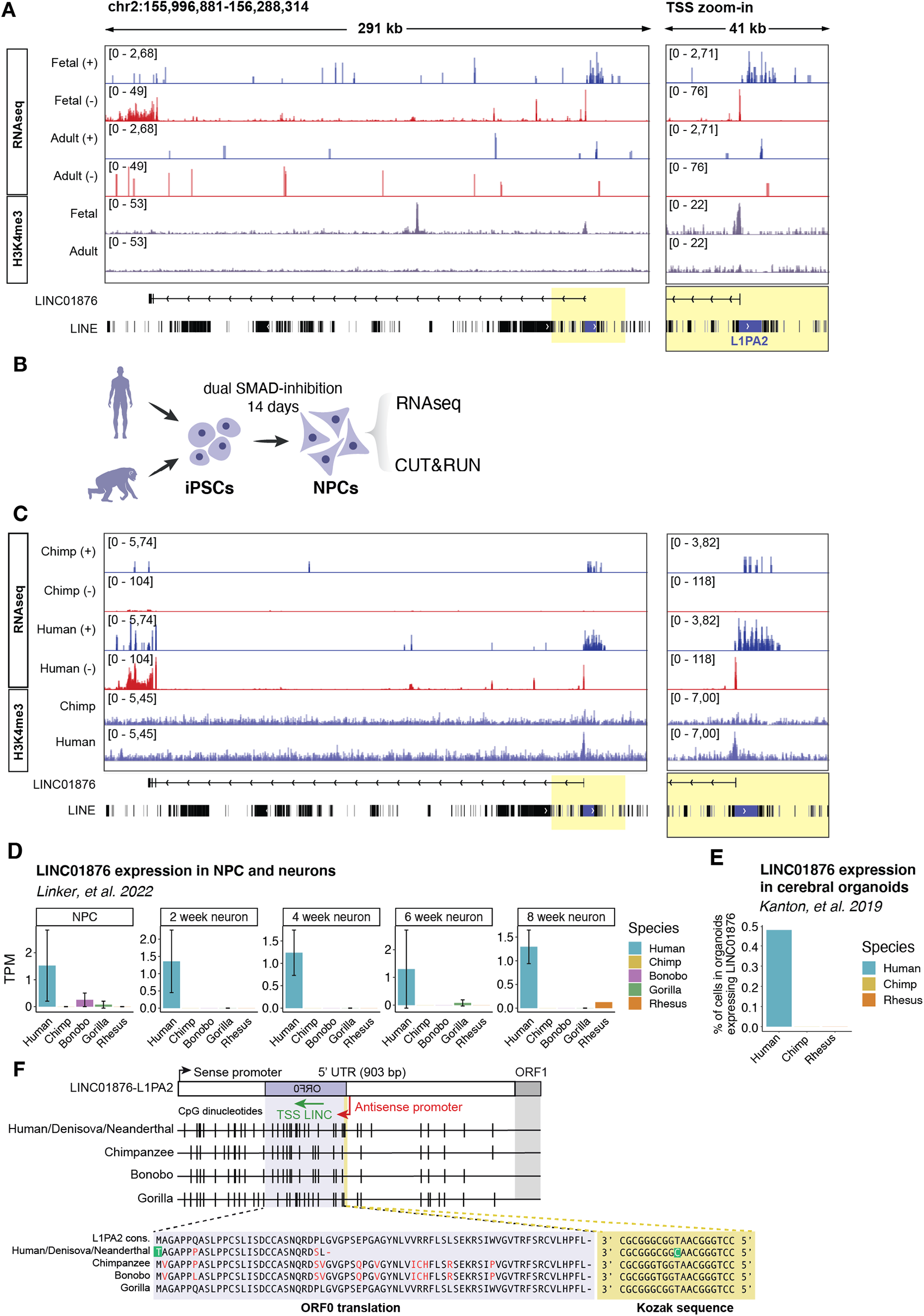
The L1-lncRNA LINC01876 is a human-specific transcript. A) Genome browser tracks showing RNAseq and H3K4me3 signal (bottom panel, in purple) over L1-lncRNA in fetal and adult samples. RNAseq signal (RPKM) split by strand (blue = forward; red = reverse). The right panel shows a zoom-in into the transcription starting site (highlighted in yellow). B) Experimental approach for fbNPCs human and chimpanzee comparison. C) Genome browser tracks showing RNAseq and H3K4me3 signal (bottom panel, in purple) over L1-lncRNA in human and chimpanzee fbNPCs. RNAseq signal (RPKM) split by strand (blue = forward; red = reverse). The right panel shows a zoom-in into the transcription starting site (highlighted in yellow). D) LINC01876 (L1-lncRNA) expression (TPM) from bulk RNAseq of human, chimpanzee, bonobo, gorilla, and macaque rhesus NPCs from Linker, et al. 2022. E) Percentage of cells expressing LINC01876 (L1-lncRNA) in human, chimpanzee, and macaque rhesus cerebral organoids from Kanton, et al. 2019. F) Multiple sequence alignment of the L1-lncRNA L1PA2 ORF0 (purple highlight) in different primates, and their Kozak sequence (yellow highlight). The transcription start site of the L1-lncRNA is indicated in orange.

### L1-lncRNA LINC01876 is a human-specific transcript

L1-derived RNAs have the potential to contribute to primate- and human speciation since they originate from the integration of new DNA sequences into our genome. To investigate the evolutionary conservation of the L1-lncRNA LINC01876 we analyzed our previously published dataset from iPSC-derived human and chimpanzee forebrain NPCs (fbNPCs) (Figure 5B) ^46^. We found the L1-lncRNA was highly expressed in human fbNPCs, as supported by both RNA-seq and H3K4me3 CUT&RUN data (Figure 5C). We were not able to detect L1-lncRNA expression in chimpanzee fbNPCs. We verified the human-specific expression of this L1-lncRNA in previously published human, chimpanzee, bonobo, gorilla and macaque RNA-seq data from NPCs and immature neurons ^47^ and snRNA-seq from human, chimpanzee and macaque cerebral organoids ^48^ (Figure 5D&E). The L1-lncRNA was consistently expressed in human NPCs, immature neurons and organoids but not in cultures obtained from other primates. Thus, the L1-lncRNA LINC01876 appears to be a human-specific transcript that is expressed during brain development.

We performed a multiple sequence alignment of the genomic region to investigate the evolutionary timepoint in which the L1PA2 was inserted into the ancestral primate genome. We found that the L1PA2 insertion site is present – and identical – in human, chimpanzee, bonobo, and gorilla, but not in orangutan, macaque or other lower species (Figure 5F). Thus, this L1PA2 insertion can be estimated to have occurred around 10-20 million years ago. To explain how the L1PA2 element drives the expression of L1-lncRNA in humans, but not in other species, we focused on its promoter region. In intact young L1s, the antisense promoter drives the expression of a small L1-peptide, ORF0 ^25^ (Figure 1G). When comparing the antisense promoter sequences of the L1PA2-insertion between humans, chimpanzees, bonobos and gorillas, we noticed a missense mutation (A451G) in the Kozak sequence of the ORF0 in humans (Figure 5F). This mutation was located at the start codon resulting in a methionine to threonine (M1T) change disabling translation of the ORF0 in humans ^25^. The ORF0 was still intact in chimpanzees, bonobos and gorillas. Denisova and Neanderthal genomes both displayed the human variant suggesting that the nucleotide change occurred before the split of archaic human species (Figure 5F). This analysis indicated that it is possible that the L1-lncRNA promoter may be silenced by DNA methylation or other repressive factors in non-human primates due to the expression and translation of an ORF0-fusion-transcript. The L1-lncRNA LINC01876 might escape silencing in humans as ORF0 is not translated, although the underlying mechanisms remain to be investigated.

### CRISPRi-mediated silencing of the L1-lncRNA reveals an important role in neural differentiation

To investigate the functional relevance of the L1-lncRNA LINC01876, we set up a CRISPRi strategy to silence LINC01876 expression. We designed 2 distinct guide RNAs (gRNA) to target unique genomic locations in the vicinity of the TSS and co-expressed these with a KRAB transcriptional repressor domain fused to catalytically dead Cas9 (KRAB-dCas9) (Figure 6A). Lentiviral transduction of human iPSCs resulted in efficient, almost complete silencing of LINC01876 upon differentiation to fbNPCs (Figure 6B, Supplemental Figure 4A) but there was no difference in differentiation capacity or expression of cell-fate markers compared to controls (Figure 6C and Supplemental Figure 4B). We also found no evidence that the expression other L1 loci was affected by the CRISPRi approach demonstrating the specificity of the silencing to the LINC01876 locus (Supplemental Figure 4C-D). The subsequently obtained results using the two different gRNAs were indistinguishable and thus results were pooled.

**Figure 6.**
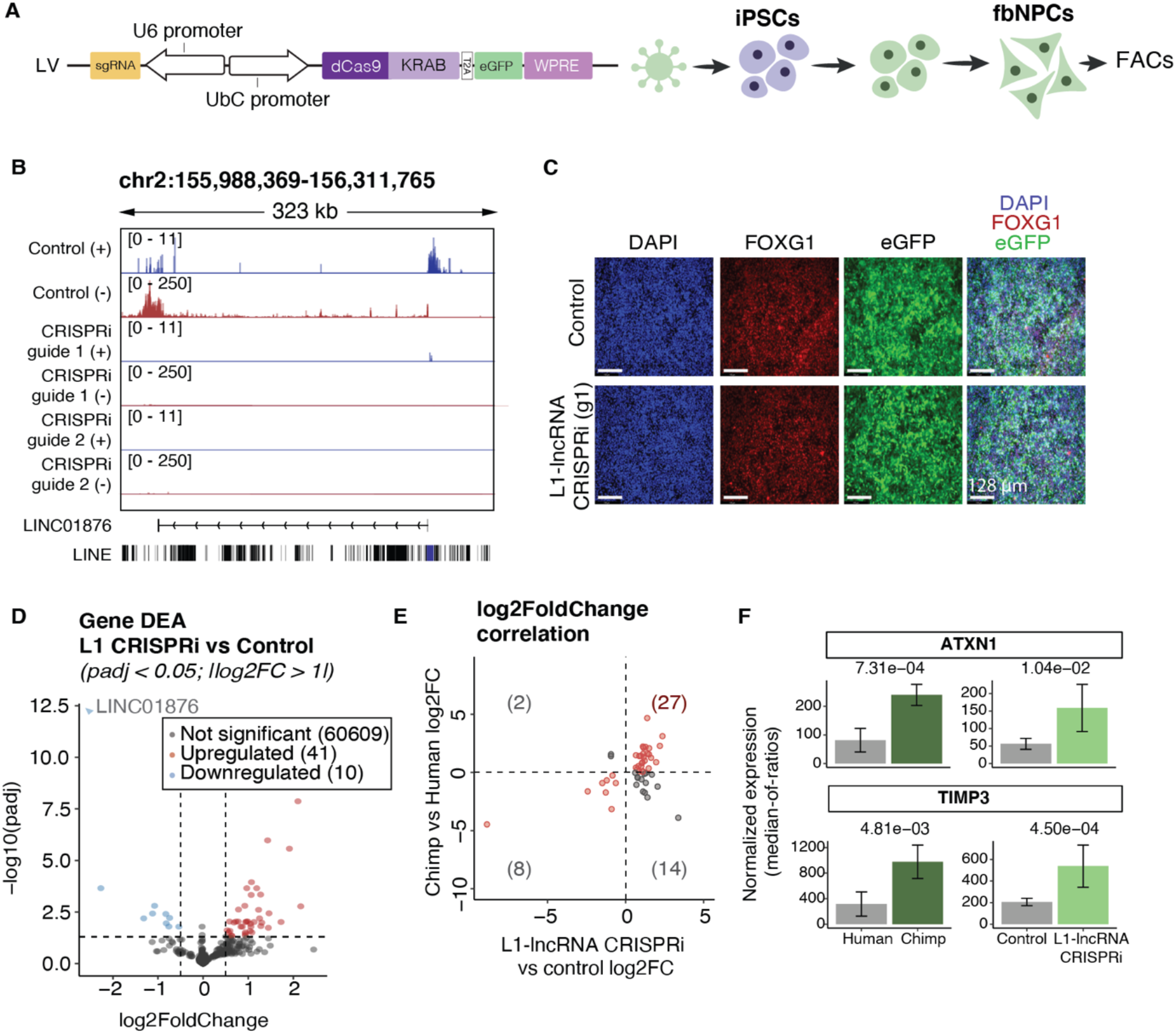
CRISPRi-silencing of the L1-lncRNA in human fbNPCs. A) CRISPRi construct and schematic of the L1-lncRNA CRISPRi in fbNPCs. B) Genome browser tracks showing the expression over L1-lncRNA in control (LacZ) and L1-lncRNA CRISPRi. RNAseq signal (RPKM) split by strand (blue = forward; red = reverse). C) Immunohistochemistry of forebrain (red = FOXG1), and nuclear (blue = DAPI) markers. eGFP showing transfected cells (green) (white scale bar 128 μm). D) Volcano plot showing differential gene expression results (DESeq2). Significantly up and downregulated genes are highlighted in red and blue respectively (log2FoldChange > 1; padj < 0.05). E) log2FoldChange of the significantly up or downregulated genes upon L1-lncRNA CRISPRi (as highlighted in D) in the two datasets (L1-lncRNA CRISPRi vs control, and human vs chimp). Genes up or downregulated in both datasets are highlighted in red (first and third quadrants). F) Normalized expression (median of ratios; DESeq2) of two example genes upregulated in both datasets.

We performed RNA-seq on LINC01876-CRISPRi fbNPCs and analyzed the transcriptome for alterations in gene expression. We found 41 significantly upregulated genes and 10 downregulated genes (DESeq2; padj < 0.05, log2FoldChange > 1) (Figure 6D). As lncRNAs can act in *cis* or *trans* ^18^ we scrutinized chromosome 2 to determine whether the differentially expressed genes were located near to the lncRNA, which would indicate a *cis* function. We found no obvious evidence suggesting that genes in the vicinity of the L1-lncRNA on chromosome 2 were affected by the CRISPRi indicating that the L1-lncRNA could act *in trans* (Supplementary Figure 4E).

We noted that many of the differentially expressed genes when comparing L1-lncRNA-fbNPCs to control-fbNPCs were also differentially expressed when comparing human and chimpanzee fbNPCs ^46^. 27 out of the 41 upregulated genes upon L1-lncRNA CRISPRi were more highly expressed in chimpanzee fbNPCs upon L1-lncRNA CRISPRi and 8 of the 10 downregulated genes after L1-lncRNA CRISPRi were expressed at lower levels in chimpanzee fbNPCs (Fig 5E). Thus, the L1-lncRNA appeared to influence the expression of several genes that distinguish the human and chimpanzee transcriptome in neural progenitors. Notably, some of these differentially expressed genes play important roles in the human brain such as Ataxin1 (ATXN1), which is mutated in spinocerebellar ataxia ^49^ and Tissue Inhibitor Of Metalloproteinases 3 (TIMP3), which is an inhibitor of the matrix metalloproteinases that has been linked to neurodegenerative disorders (Figure 5F) ^50^.

### L1-lncRNA LINC01876 contributes to developmental timing in cerebral organoids

To investigate the functional role of the L1-lncRNA in human brain development, we generated L1-lncRNA-CRISPRi cerebral organoids. This model allows for the study of human-specific developmental processes in 3D ^51^ (Figure 7A). We found that L1-lncRNA-CRISPRi silencing did not impair the organoid formation and the resulting organoids displayed characteristic neural rosettes after 30 days of growth, as visualized with Pax6/ZO1 staining (Figure 7B). Quantification of organoid size throughout differentiation revealed that L1-lncRNA-CRISPRi organoids were reproducibly smaller than control organoids (Figure 7C, Supplemental Figure 5B). This difference appeared after two weeks of growth and was sustained up until 1 month, which was the last time point quantified (Figure 7C). The results were independently reproduced using two different gRNAs.

**Figure 7.**
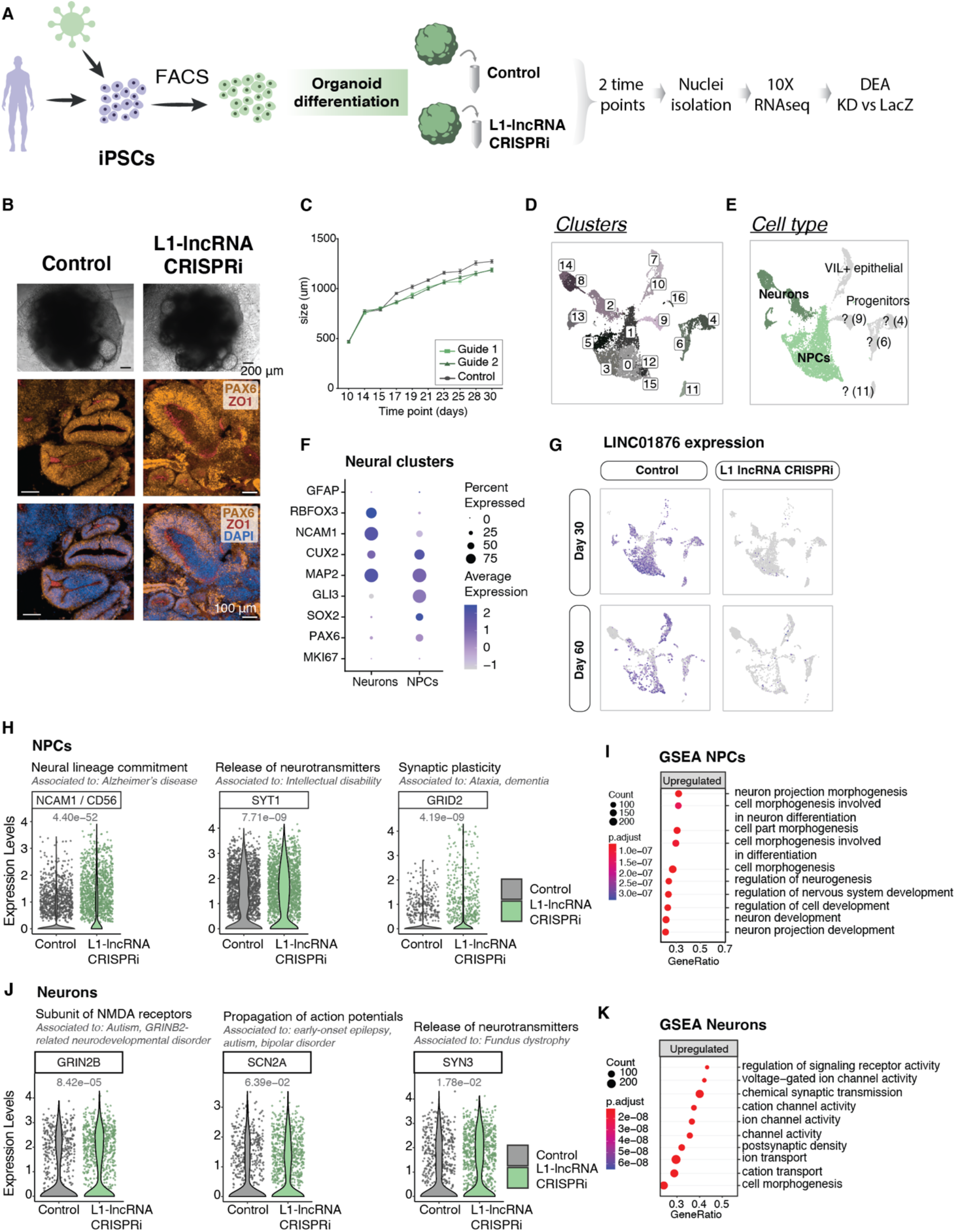
Silencing of L1-lncRNA in cerebral organoids indicates it has a role in developmental timing. A) Schematic of experimental design for organoid differentiation, L1-lncRNA CRISPRi, and sequencing. B) Bright-field pictures of iPSC-derived cerebral organoids (top, black scale bar 200 μm). Immunohistochemistry of PAX6 (orange), ZO1 (red) and DAPI (blue) (bottom, white scale bar 100 μm). C) Quantification of organoid diameter between days 10 and 30 (n=20-30 organoids per time point) D) Single nuclei RNAseq: UMAP colored by cluster E) UMAP colored by identified cell types. Neuronal-like clusters colored in two shades of green, uncharacterized clusters or progenitor-like cells colored in grey. F) Dot plot showing expression of neuronal and neuronal progenitor markers in the NPC and neuronal clusters. G) UMAP showing expression of L1-lncRNA. H) Selected examples of significantly upregulated genes in L1-lncRNA CRISPRi NPCs (FindMarkers from Seurat; padj < 0.05). I) Selected upregulated terms of the gene set enrichment analysis (GSEA) over NPCs (gseGO; padj < 0.05). J) Selected examples of significantly upregulated genes in L1-lncRNA CRISPRi Neurons (FindMarkers from Seurat; padj < 0.05). K) Selected upregulated terms of GSEA over Neurons (gseGO; padj < 0.05).

To further evaluate the long-term molecular consequences of L1-lncRNA inhibition on human brain development, we analyzed organoids at 1 and 2 months of growth using snRNA-seq. High-quality data were generated from a total of 11,669 cells, including 6,099 from L1-lncRNA-CRISPRi organoids (2 gRNAs, in total 45 organoids) and 5,570 from control organoids (lacZ-gRNA, in total 25 organoids). We performed an unbiased clustering analysis to identify and quantify the different cell types present in the organoids. 17 separate clusters were identified (Figure 7D), including cerebral cells of different stages of maturation, such as NPCs and newborn neurons (Figure 7E-F). All of the clusters contained cells from both 1 and 2 months, and we found no apparent difference in the contribution to the different clusters by L1-lncRNA-CRISPRi organoids, suggesting that the L1-lncRNA LINC01876 does not influence developmental fate during early human brain development (Supplemental Figure 5A).

Next, we analyzed the transcriptional difference between control and L1-lncRNA-CRISPRi organoids. We confirmed the transcriptional silencing of L1-lncRNA in all cell populations at both time points (Figure 7G). Notably, in ctrl-organoids the L1-lncRNA was expressed in NPCs but not in neurons, demonstrating that the 3D-system is able to replicate an appropriate developmentally regulated expression pattern of this L1-derived transcript (Figure 7G). We found that in the NPC population, genes linked to neuronal differentiation, such as NCAM1, SYT1, and GRID2, were upregulated in L1-lncRNA-CRISPRi organoids (Figure 7H). An unbiased gene set enrichment analysis (GSEA) of the upregulated genes in NPCs was significantly enriched in gene ontology (GO) terms linked to neuronal differentiation (Figure 7I). In line with this, we found that in newborn neurons, genes linked to mature neuronal functions, such as GRIN2B, SCN2A and SYN3, were upregulated in L1-lncRNA-CRISPRi organoids (Figure 7J) and GSEA confirmed enrichment of upregulated genes linked to neuronal maturation (Figure 7K). These results indicate that NPCs and neurons present in organoids that lack the L1-lncRNA LINC01876 display a more mature transcriptional profile than those found in control cerebral organoids.

Taken together, these results demonstrate that silencing of the L1-lncRNA LINC01876 results in organoids that contain the same cell types as control organoids, suggesting that the L1-lncRNA does not influence developmental fate. However, we found that the L1-lncRNA organoids were smaller during early differentiation and displayed transcriptome changes in line with more mature NPCs and neurons. These observations are in line with a role for the L1-lncRNA in developmental timing since L1-lncRNA-CRISPRi organoids appear to differentiate more quickly.

## Discussion

L1 mobilization represents a threat to human genomic integrity, and it has therefore been assumed that L1 expression is silenced in somatic human tissues. However, the abundance and repetitive nature of L1s make their transcription difficult to precisely estimate ^34^. Previous studies have, based on retrotransposition activity, indirectly indicated that L1s may be expressed in the brain, but the available data is conflicting, and a clear consensus has not been established ^27–33^. Therefore, several open questions remain: are L1s expressed in the human brain and in what cell types? Are L1s developmentally regulated? Does L1-derived transcripts contribute to brain functions? In this study we resolve many of these issues through the use of a careful multi-omics anaylsis of human tissue, combined with a customised bioinformatic pipelines. We found that L1s are highly expressed in the developing human brain and in neurons in the adult human brain.

Our data demonstrates that the expression of L1s in the developing and adult human brain is largely limited to evolutionarily young, primate-specific L1s, primarily subclasses found only in hominoids. The lack of expression of more ancient L1s is likely explained by the higher burden of deletions, mutations and genomic rearrangements of old TEs that reduce their capacity to be transcribed. Importantly, a strand-specific analysis of full-length elements that contain an intact 5’ promoter revealed that the RNA-seq signal was present in sense to the L1s. We thereby confirmed that hundreds of different L1 loci are expressed, and that the L1 signal is not transcriptional noise – but rather that the L1 promoter drives expression. This strongly suggests that the signal is not the result of passive expression in which the L1 sequence is incorporated into another transcript ^34^. We confirmed this with two orthogonal approaches: by performing long-read RNA-seq analysis to identify L1 transcripts that initiate in the L1 5’ UTR, and by H3K4me3-profling to identify L1 promoters active in the human brain – benefiting from the fact that the signal of this histone modification spreads to the flanking (and thus unique) genomic context. We thus found *bona fide* evidence that full-length L1s are expressed in both the developing and adult human brain. However, we acknowledge that with our approach we miss the expression of polymorphic L1s not present in the reference genome. Future studies using individual-matched RNA-seq and long-read genome data will be crucial to investigate if L1s individualizes the neuronal transcriptome.

From our analysis, it is evident that not all L1 loci are expressed in the brain, but rather a small subset. Our data also indicate that the L1 integration site is important and that the presence of highly active nearby gene promoters or other regulatory elements is key for L1 expression. Thus, the activity of the surrounding genome is one parameter that is important for how this subset of L1s escape silencing. In this respect, our results are similar to what have previously been found in cancer cell lines ^45^. In addition, single nucleotide variants or small deletions in regulatory regions of individual L1 integrant could result in the avoidance of recruiting silencing factors. A previous study indicated that a subset of young L1s that have lost a YY1-binding site in the promoter avoids silencing in the brain in a DNA methylation-dependent manner ^32^. However, in our dataset we found L1s both with and without the YY1 binding site to be expressed (Supplemental Figure 6A-B). Another interesting aspect of our data is that L1HS elements appears to be globally silenced in brain development. This indicates that L1HS elements are controlled by unique, specialized mechanisms during brain development, likely to avoid abundant retrotransposition events in proliferating cell populations. The nature of this mechanism remains unknown, but it will be interesting to investigate further to understand how the human brain avoids waves of retrotransposition events during early development and what the consequences are if this mechanism fails.

The fact that many L1 promoters are active in the human brain demonstrate that L1s are a rich source of genetic sequences that provides a primate-specific layer of transcriptional complexity. Our data indicates that L1s influence the expression of protein-coding genes and non-coding transcripts in the human brain through several mechanisms, including acting as alternative promoters, enhancer elements or by altering 5’ and 3’UTRs. In addition, there is the possibility that L1-derived peptides or fusion peptides play important functional roles ^52^. One example of an L1-derived non-coding transcript that we identified is LINC01876, a L1-lncRNA that exploits the antisense promoter of a L1PA2 element that is transcriptionally active in human brain development. In the LINC01876 promoter, the first amino acid of ORF0 is specifically mutated in humans and the subsequent loss of ORF0 coding capacity correlates with the appearance of the L1-lncRNA. It is possible that this single nucleotide variant, at a key position for the L1-lifecycle, enables the escape of DNA methylation-mediated silencing resulting in transcription of the lncRNA.

Our loss-of-function studies of the L1-lncRNA LINC01876 indicate that it plays an important role in regulating developmental timing during human brain development. LINC01876 is a previously uncharacterized lncRNA, but we have noted that in the promoter region of LINC01876 there is a SNP that is linked to major depressive disorders ^53^. Our data demonstrates that organoids in which LINC01876 expression was silenced were smaller in size and displayed NPCs and neurons with a more mature transcriptome than control counterparts. These findings are reminiscent of previously observed differences when comparing human cerebral organoids to those derived from non-human great apes ^48, 54, 55^. Thus, our data provides direct experimental evidence as to how an L1 insertion has contributed to the evolution of the human brain and provide novel links between L1s and the genetics of neuropsychiatric disorders that will be interesting to study in detail in the future.

In summary, our results illustrate how L1s provide a layer of transcriptional complexity in the brain and provides evidence for the contribution of one such example to the evolution of the human brain. Our results establish L1s as powerful genetic elements with relevance in human brain function. It has been estimated that a new L1 germline insertion occurs in every 50-200 human births ^9, 40^. This extensive L1 mobilization in the human population has resulted in hundreds of unfixed polymorphic L1-insertions in each human genome ^9, 56^. Since L1s are highly polymorphic within the human population, the prevalence of certain L1 copies or SNPs and structural variants in fixed L1s in the genome is therefore likely to influence the etiology of brain disorders. Thus, L1s represent a set of genetic material that have been important in the evolution of our brain and likely contribute to many important gene regulatory and transcriptional networks in the human brain. L1s should no longer be neglected, and these sequences need to be included in future investigations of the underlying genetic causes of human brain disorders.

### Methods iPSCs

Human induced pluripotent stem cell (iPSC) line generated by mRNA transfection was used: RBRC-HPS0328 606A1, hereafter referred to as HS1 (Riken, RRID:CVCL_DQ11). The iPSC line was maintained as previously described ^46, 57, 58^. Briefly, the iPSC lines were maintained on LN521 (0.7 µg/cm2; Biolamina) coated Nunc multi-dishes in iPS media (StemMACS iPS-Brew XF and 0.5% penicillin/streptomycin; GIBCO) and were passaged 1:2-1:6 every 2-4 days once 70-90% confluency was reached. The media was changed daily and 10 μM Y27632 (Rock inhibitor, Miltenyi) was added when cells were passaged.

### Forebrain Neural Progenitor Cells (fbNPCs)

iPSCs were differentiated into forebrain neural progenitors (fbNPCs) as previously described ^46, 57^. Upon dissociation at 70-90% confluency, the cells were plated on LN111 (1.14µg/cm2; Biolamina) coated Nunc multidishes at a density of 10.000 cells/cm2 and grown in N2 medium (1:1 DMEM/F-12 (21331020; GIBCO) and Neurobasal (21103049; GIBCO) supplemented with 1% N2 (GIBCO), 2 mM L-glutamine (GIBCO), and 0.2% penicillin/streptomycin). 10 μM SB431542 (Axon) and 100 ng/ml noggin (Miltenyi) were given for dual SMAD inhibition. The media was changed every 2-3 days. On day 9, N2 media without dual SMAD inhibitors were used. On day 11, cells were dissociated and replated on LN111 coated Nunc multidishes at a density of 800.000 cells/cm2 in B27 medium (Neurobasal supplemented with 1% B27 without vitamin A (GIBCO), 2 mM L-glutamine and 0.2% penicillin/streptomycin Y27632 (10 μM), BDNF (20 ng/ml; R&D), and L-ascorbic acid (0.2 mM; Sigma). Cells were kept in the same media until day 14 when cells were harvested for downstream analysis.

### CRISPRi

To silence the expression of LINC01876 in iPSCs, we adapted the protocol detailed in ^46^. Single guide sequences were designed to recognize DNA regions near the transcription start site (TSS) according to the GPP Portal (Broad Institute). The guide sequences were inserted into a deadCas9-KRAB-T2A-GFP lentiviral backbone, pLV hU6-sgRNA hUbC-dCas9-KRAB-T2a-GFP, a gift from Charles Gersbach (Addgene plasmid #71237 RRID:Addgene_71237), using annealed oligos and the BsmBI cloning site. Lentivirus was produced as described below and iPSCs transfected with MOI 10 of LacZ and LINC01876-targeting guide RNA. Guide efficiency was validated using standard quantitative real-time RT-PCR techniques

LINC01876 guide 2: ACGAGATTATAAGCCGCACC

LINC01876 guide 3: AGGGGCGCCCGCCGTTGCCC

LacZ: TGCGAATACGCCCACGCGAT

GFP+ cell isolation of fbNPCs: At day 14 cells were detached with accutase, resuspended in B27 media containing RY27632 (10 μM) and Draq7 (1:1000, BD Bioscience) and strained with a 70μm (BD Bioscience) filter. Gating parameters were determined by side and forward scatter to eliminate debris and aggregated cells. The GFP-positive gates were set using untransduced fbNPCs. The sorting gates and strategies were validated via reanalysis of sorted cells (> 95% purity cut-off). 200.000 GFP-positive/Draq7-negative cells were collected per sample, spun down at 400g for 5 min and snap-frozen on dry ice. Cell pellets were kept at −80°C until RNA was isolated.

GFP+ cell isolation of transduced iPSCs: 7 days post-transduction, cells were detached with accutase, resuspended in iPS media containing RY27632 (10 μM) and Draq7 (1:1000) and strained with a 70μm filter. Gating parameters were determined by side and forward scatter to eliminate debris and aggregated cells. The GFP-positive gates were set using untransduced iPSCs. The sorting gates and strategies were validated via reanalysis of sorted cells (> 95% purity cut-off). 200.000 GFP-positive/Draq7-negative cells were collected per sample, spun down at 400g for 5 min and resuspended in iPS media containing RY27632 (10 μM) and expanded as described above and frozen down for further use.

Detailed protocol can be found at DOI: dx.doi.org/10.17504/protocols.io.yxmvm25n9g3p/v1.

### Lentiviral production

Lentiviral vectors were produced according to Zufferey et al ^59^ and were titered by qRT-PCR. Briefly, HEK293T cells (RRID:CVCL_0063) were grown to a confluency of 70 – 90% for lentiviral production. Third-generation packaging and envelope vectors (pMDL (Addgene #12251), psRev (Addgene #12253), and pMD2G (Addgene #12259)) together with Polyethyleneimine (PEI Polysciences PN 23966, in DPBS (GIBCO) were used in conjunction with the lentiviral plasmids previously generated. The lentivirus was harvested two days after transfection. The media was collected, filtered and centrifuged at 25.000g for 1.5 hours at 4°C. The supernatant was removed from the tubes and the virus was resuspended in DPBS and left at 4°C. The resulting lentivirus was aliquoted and stored at −80°C.

### qRT-PCR

Total RNA was first extracted using the miniRNeasy kit (QIAGEN). cDNA was generated using the Maxima First Strand cDNA Synthesis Kit (Thermo Scientific). Quantitative qPCR was performed using SYBR Green I master (Roche) on a LightCycler 480 (Roche). The 2-ΔΔCt method was used to normalize expression to control, relative to GAPDH and B-ACTIN as described previously ^60^.

### List of Primers used

Gene Primer sequence (5’ to 3’)

LINC01876 Fw AATCCGTGCCAGCAGTAAGT Rev GGACCTCTTCAAGTCCCAGG ACTB Fw CCTTGCACA TGCCGGAG Rev GCACAGAGCCTCGCCTT GAPDH Fw TTGAGGTCAARGAAGGGGTC Rev GAAGGTGAAGGTCGGAGTCA

### Human cerebral organoid culture

To generate the human cerebral-like organoids we followed the protocol detailed in ^46^. We used three hIPSC6-derived lines obtained by transduction and FACS sorting as described above: one control line (guide against LacZ) and two LINC01876 CRISPRi KD lines (guide 2 and guide 3). Briefly, 8000 cells/well were plated in a 96-wells plate (Costar, Ultra Low Attachment, round bottom; REF 7007) with 250 μL of mTeSR1 (StemCell Technologies, Inc.) and RY27632 10 μM. This is considered day −5 of the differentiation of the iPSCs-derived hFB organoids. On days −3 and −1 the medium was changed (150 μL and 200 μL of mTeSR1, respectively). At day 0 the cells are fed with Neural Induction Medium (NIM; DMEM/F12 media, N2 Supplement (1:100), L-Glutamine (2mM), Penicillin/Streptomycin (1:500), Non-Essential Amino acids (1:100) and Heparin (2ug/ml).) enriched with 3% KSR. On days 2, 4, and 6, the organoids were fed with NIM with no added KSR.

On day 8 the organoids were embedded in 30-50 μL of Matrigel (Corning) and incubated at 37°C for 25 minutes to allow the Matrigel to solidify. The organoids were then transferred in Corning REF 3471 6-well plates with flat bottoms containing 4 ml/well of Cortical Differentiation Medium (CDM; F12 Media (-Glut) (48.5%), Neurobasal (48.5%), N2 Supplement (1:200), B27 Supplement (-Vit.A, 1:100), L-Glutamine (2mM), Penicillin/Streptomycin (1:500), Non-Essential Amino acids (1:200), Beta MercaptoEtOH (50uM) and Insulin (2.5 ug/mL)).

On days 10 and 12 of the differentiation, the media was changed exchanging 3 ml/well for 3 mL of fresh CDM. On days 15, 17, 19, 21 and 23, ∼4 mL of the medium was replaced with 4 mL of Improved Differentiation Medium + A (IDM, F12 Media (-Glut) (48.5%), Neurobasal (48.5%), N2 Supplement (1:200), B27 Supplement (+Vit.A, 1:50), L-Glutamine (2mM), Penicillin/Streptomycin (1:500), Non-Essential Amino acids (1:200), Beta MercaptoEtOH (50uM), Insulin (2.5 ug/mL) and Ascorbic Acid (400uM)). From day 25, the media was changed every 3 days with 3-4 mL of Cortical Terminal Differentiation Medium (CTDM, F12 Media (-Glut) (48.5%), Neurobasal (48.5%), N2 Supplement (1:200), B27 Supplement (+Vit.A) – (1:50) 800uL, L-Glutamine (2mM), Penicillin/Streptomycin (1:500), Non-Essential Amino acids (1:200), Beta MercaptoEtOH (50uM), Insulin (2.5 ug/mL) and Ascorbic Acid (400uM), BDNF (10ng/uL), cAMP (200uM), GDNF (10ng/uL)).

All the diameter measurements of the organoids were taken with the Measure tool from Image J (RRID:SCR_003070). The chosen measuring unit was μm.

Detailed can be found at DOI: dx.doi.org/10.17504/protocols.io.e6nvwjo27lmk/v1.

### Immunocytochemistry

The cells were washed three times with DPBS and fixed for 10 minutes with 4% paraformaldehyde (Merck Millipore), followed by three more rinses with DPBS. The fixed cells were then blocked for 60 minutes in a blocking solution of KPBS with 0.25% Triton X-100 (Fisher Scientific) and 5% donkey serum at room temperature.

The primary antibody (rabbit anti-FOXG1 (Abcam; RRID: AB_732415), 1:50) was added to the blocking solution and incubated overnight at room temperature. Subsequently, the cells were washed three times with KPBS. The secondary antibody (donkey anti-rabbit Cy3 (Jackson ImmunoResearch Labs Cat# 711165152; RRID:AB_2307443), 1:200) was added with DAPI (1:1000, Sigma-Aldrich) to the blocking solution and incubated at room temperature for one hour, followed by 2-3 rinses with KPBS. The cells were visualized on a Leica microscope (model DMI6000 B).

Detail protocol can be found at DOI: dx.doi.org/10.17504/protocols.io.5qpvor7pdv4o/v1.

### Immunohistochemistry

Organoids were fixed in 4% paraformaldehyde for 2 hours at room temperature. They were subsequently washed three times with KPBS and left in a 1:1 30% sucrose solution and OCT (HistoLab, Cat# 45830) mixture overnight at 4°C. Organoids were then transferred to a cryomold containing OCT, frozen on dry ice and stored at -80°C in freezer bags.

Prior to staining, organoids were sectioned on a cryostat at -20 °C at a thickness of 20 μM and placed onto Superfrost plus microscope slides. They were then washed 3x with KPBS for 5 minutes and subsequently blocked and permeabilized in 0.1% Triton X-100 and 5% normal donkey serum in KPBS for one hour at room temperature. The primary antibody (rabbit anti-PAX6 (Biolegend, Cat# 901301, RRID:AB_2565003) 1:300 dilution, and rat anti-ZO1 (Novus, Cat# NB110-68140, RRID:AB_1111431) 1:300 dilution) was added to the blocking solution and incubated overnight at room temperature. Subsequently, the sections were washed three times with KPBS. The secondary antibody (donkey anti-rabbit Cy3 (Jackson ImmunoResearch Labs, Cat# 711165152, RRID:AB_2307443); 1:200 and donkey anti-rat Cy5 (Jackson ImmunoResearch Labs, Cat# 712175153; RRID: AB_2340672), 1:200) was added with DAPI (1:1000; Sigma-Aldrich) to the blocking solution and incubated at room temperature for one hour, followed by 2-3 rinses with KPBS. Sections were imaged using Operetta CLS (PerkinElmer).

Detail protocol can be found at DOI: dx.doi.org/10.17504/protocols.io.n92ldp22nl5b/v1.

### Single nuclei isolation

The nuclei isolation from the embryonic brain tissue and organoids was performed according to ^36^. In brief, the tissue and organoids were thawed and dissociated in ice-cold lysis buffer (0.32 M sucrose, 5 mM CaCl2, 3 mM MgAc, 0.1 mM Na2EDTA, 10 mM Tris-HCl, pH 8.0, 1 mM DTT) using a 1 ml tissue douncer (Wheaton). The homogenate was carefully layered on top of a sucrose cushion (1.8 M sucrose, 3 mM MgAc, 10 mM Tris-HCl, pH 8.0, and 1 mM DTT) before centrifugation at 30,000 × g for 2 hours and 15 min. Pelleted nuclei were softened for 10 min in 100 ml of nuclear storage buffer (15% sucrose, 10 mM Tris-HCl, pH 7.2, 70 mM KCl, and 2 mM MgCl2) before being resuspended in 300 ml of dilution buffer (10 mM Tris-HCl, pH 7.2, 70 mM KCl, and 2 mM MgCl2) and run through a cell strainer (70 mm). Cells were run through the FACS (FACS Aria, BD Biosciences) at 4°C at a low flow rate using a 100 mm nozzle (reanalysis showed >99% purity). Nuclei intended for bulk RNA sequencing were pelleted at 1,300 x g for 15 min.

Detail protocol can be found DOI: dx.doi.org/10.17504/protocols.io.5jyl8j678g2w/v1.

### 3’ and 5’ single nuclei sequencing

Nuclei or cells intended for single cell/nuclei RNA sequencing (8,500 nuclei/cells per sample) were directly loaded onto the Chromium Next GEM Chip G or Chromium Next GEM Chip K Single Cell Kit along with the reverse transcription mastermix following the manufacturer’s protocol for the Chromium Next GEM single cell 3’ kit (10X Genomics, PN-1000268) or Chromium Next GEM Single Cell 5’ Kit (10x Genomics, PN-1000263) respectively, to generate single-cell gel beads in emulsion. cDNA amplification was done as per the guidelines from 10x Genomics using 13 cycles of amplification for 3’ and 15 cycles of amplification for 5’ libraries. Sequencing libraries were generated with unique dual indices (TT set A) and pooled for sequencing on a Novaseq6000 using a 100-cycle kit and 28-10-10-90 reads.

#### Single cell/nuclei RNAseq analysis

##### Gene quantification

The raw base calls were demultiplexed and converted to sample-specific fastq files using 10x Genomics Cell Ranger mkfastq (version 3.1.0; RRID:SCR_017344) ^61^. Cell Ranger count was run with default settings, using an mRNA reference for single-cell samples and a pre-mRNA reference (generated using 10x Genomics Cell Ranger 3.1.0 guidelines) for single nuclei samples.

To produce velocity plots, loom files were generated using velocyto ^43^ (version 0.17.17; RRID:SCR_018167) run10x in default parameters, masking for TEs (same GTF file as input for TEtranscripts; see method section Bulk RNA sequencing analysis <u>TE subfamily quantification</u>) and gencode version 36 as guide for features. Plots were generated using velocyto.R (see github under src/analysis/ fetal_velocity.Rmd).

#### Clustering

Samples were analysed using Seurat (version 3.1.5; RRID:SCR_007322) ^62^. For each sample, cells were filtered out if the percentage of mitochondrial content was over 10% (perc_mitochondrial). For adult samples, cells were discarded if the number of detected features (nFeature_RNA) was higher than 2 standard deviations over the mean in the sample (to avoid keeping doublets), or lower than a standard deviation below the mean in the sample (to avoid low quality cells). For fetal samples, cells were discarded if the number of detected features was higher than 2 standard deviations over the mean in the sample, or lower than 2,000 features detected. Counts were normalized using the Centered Log Ratio (CLR) transformation (Seurat::NormalizeData) and clusters were found with a resolution 0.5 (Seurat::FindClusters).

##### TE quantification

We used an in-house pseudo-bulk approach to process single nuclei RNAseq data to quantify TE expression per cluster, similar to what has been previously described ^36^. All clustering, normalization and merging of samples were performed using the contained scripts of get_clusters.R (get_custers() from the Sample class) and merge_samples.R (merge_samples() from the Experiment class) of trusTEr (version 0.1.1; doi:10.5281/zenodo.7589548). Documentation of the pipeline can be found at https://raquelgarza.github.io/truster/.

The main functionality of trusTEr is to create collections of reads to remap and quantify TE subfamilies or elements per group of cells. The function tsv_to_bam() backtraces cells barcodes to Cell Ranger’s output BAM file. tsv_to_bam() runs using subset-bam from 10x Genomics version 1.0 (RRID:SCR_023216). As the next step of the pipeline, the function filter_UMIs() filters potential PCR duplicates in the BAM files; this step uses Pysam version 0.15.1 (RRID:SCR_021017). Next, to convert BAM to FastQ files, we used bamtofastq from 10x Genomics (version 1.2.0; RRID: SCR_023215). The remapping of the clusters was performed using STAR aligner (version 2.7.8a; RRID:SCR_004463). Quantification of TE subfamilies was done using TEcount (version 2.0.3; RRID:SCR_023208) and individual elements were quantified using featureCounts (Subread version 1.6.3; RRID:SCR_012919). The normalization step of trusTEr, to integrate with Seurat and normalize TE subfamilies’ expression, was performed using Seurat version 3.1.5 (RRID:SCR_007322).

For the purposes of this paper, we combined the samples from the same condition (all embryonic samples and all adult samples). The quantification was run twice: with all samples together, and per sample in the combined clustering. For the fetal samples, we also ran trusTEr grouping clusters per cell cycle state, for which we prepared a directory with tsv files containing the barcodes of the cells in each of the clusters of interest (e.g. cluster0_cycling.tsv, cluster0_noncycling.tsv, …) and ran the set_merge_samples_outdir function from the Experiment class to register these as cluster objects.

### Bulk RNA sequencing

Total RNA was isolated from nuclei, cell culture samples, or tissue using the RNeasy Mini Kit (Qiagen). Libraries were generated using Illumina TruSeq Stranded mRNA library prep kit (poly-A selection) and sequenced on a NextSeq500 (PE 2×150bp).

Protocol can be found at DOI: dx.doi.org/10.17504/protocols.io.36wgqjqbkvk5/v1.

#### Bulk RNA sequencing analysis

##### TE subfamily quantification

For the quantification of transposable element subfamilies, the reads were mapped using STAR aligner (version 2.6.0c; RRID:SCR_004463) ^63^ with an hg38 index and gencode version 36 as the guide GTF (--sjdbGTFfile), allowing for a maximum of 100 multi mapping loci (--outFilterMultimapNmax 100) and 200 anchors (-- winAnchorMultimapNmax). The rest of the parameters affecting the mapping were left in default as for version 2.6.0c.

The TE subfamily quantification was performed using TEcount from the TEToolkit (version 2.0.3; RRID:SCR_023208) in mode multi (--mode). Gencode annotation v36 was used as the input gene GTF (--GTF), and the provided hg38 GTF file from the author’s web server as the TE GTF (--TE) ^35^.

##### TE quantification

Reads were mapped using STAR aligner (version 2.6.0c; RRID:SCR_004463) ^63^ with an hg38 index and gencode version 30 (adult data) and 36 (fetal data) as the guide GTF (--sjdbGTFfile). To quantify only confident alignments, we allowed a single mapping locus (--outFilterMultimapNmax 1) and a ratio of mismatches to the mapped length of 0.03 (--outFilterMismatchNoverLmax).

To measure the antisense transcription over a feature, we divided the resulting BAM file into two, containing the forward and reverse transcription respectively. We used SAMtools view (version 1.9; RRID:SCR_002105) ^64^ to keep only the alignments in forward transcription, we separated alignments of the second pair mate if they mapped to the forward strand (-f 128 -F 16) and alignments of the first pair mate if they map to the reverse strand (-f 80). To keep the reverse transcription, we kept alignments of the second pair mate if they mapped to the reverse strand (-f 144) and alignments of the first pair mate if they mapped to the forward strand (-f 64 -F 16).

Both BAM files were then quantified using featureCounts from the subread package (version 1.6.3; RRID:SCR_012919) (Liao et al., 2014) forcing strandness to the features being quantified (-s 2). For consistency (and to avoid quantifying over simple repeats, small RNAs and low-complexity regions) we input the same curated hg38 GTF file provided by the TEtranscripts authors ^35^.

##### Gene quantification

Reads were mapped using STAR aligner (version 2.6.0c; RRID:SCR_004463) ^63^ with an hg38 index and gencode version 36 as the guide GTF (--sjdbGTFfile), no other parameters were modified (default values for --outFilterMultimapNmax, -- outFilterMismatchNoverLmax, and --winAnchorMultimapNmax).

Genes were quantified using featureCounts from the subread package (version 1.6.3; RRID:SCR_012919) ^65^ forcing strandness (-s 2) to quantify by gene_id (-g) from the GTF of gencode version 36.

##### Differential gene expression analysis

We performed differential expression analysis using DESeq2 (version 1.28.1; RRID:SCR_015687) ^66^ with the read count matrix from featureCounts (Subread version 1.6.3; RRID:SCR_012919) as input. Fold changes were shrunk using DESeq2:: lfcShrink.

For the produced heatmaps, counts were normalized by median-of-ratios as described by Love et al, 2014, summed with a pseudo count of 0.5 and log2 transformed.

For further detail, please refer to the Rmarkdown on the github.

##### Differential TE subfamilies expression analysis

We performed differential expression analysis using DESeq2 (version 1.28.1; RRID:SCR_015687) ^66^ with the read count matrix from TEcount (version 2.0.3; RRID:SCR_023208) ^35^ using only the TE subfamilies entries. Fold changes were shrinked using DESeq2:: lfcShrink.

Using the gene DESeq2 object (see section above) we normalized the TE subfamily counts by dividing the read count matrix by the sample distances (sizeFactor) as calculated by DESeq2 with the quantification of genes without multimapping reads (see section “Gene quantification”). For heatmap visualization, a pseudo count of 0.5 was added and log2 transformed.

##### Comparison between sense and antisense transcription over TEs

To normalize uniquely mapped read counts per strand (see section “TE quantification”), we divided the read count matrix by the sample distances (sizeFactor) as calculated by DESeq2 (version 1.28.1; RRID:SCR_015687) with the quantification of genes without multimapping reads (see section “Gene quantification”).

Each point in the boxplot (Figures 1E and 4E) refers to a sample. “Antisense” refers to counts of reverse transcription in forward features and counts from forward transcription in reverse features. “Sense” refers to counts of reverse transcription in features annotated in the reverse strand, and forward counts in features annotated in the forward strand. Boxplots were produced by summing counts of the same subfamily and strand, per sample, per the direction of transcription (e.g., all L1PA2s in the reverse strand were summed using only the counts from the reverse strand).

##### Comparing the ratio of detected elements of all L1s

Once normalized for the counts of individual elements by the gene sizeFactors (see “Comparison between sense and antisense transcription over TEs” (Figure 1E and 4E) section), we defined a “detected” element as an element with a mean >10 normalized counts in the group of samples of interest. The total number of elements is the number of elements from a particular subfamily annotated in the GTF file that was input to featureCounts (version 1.6.3; RRID:SCR_012919).

### Transcription over evolutionary young L1s elements in bulk datasets

The BED file version of TEcount’s GTF file was used to create BED files containing all L1HS, L1PA2, L1PA3, and L1PA4 elements longer than 6 kbp (full length). These BED files were then split by the strand of the element.

Using the bigwig files of the uniquely mapped BAM files, we created four matrices per dataset using the deeptools’ (version 2.5.4; RRID:SCR_016366) computeMatrix function ^67^ – one for elements annotated in the positive strand using only the bigwig files with forward transcription (transcription in sense of the element), another one for elements annotated in the reverse strand using only bigwig files with reverse transcription (transcription in sense of the element), and another two with the antisense transcription being used (e.g. elements annotated in the positive strand using reverse transcription bigwig files). We then concatenate the matrices of transcription in sense of the elements together using rbind from computeMatrixOperations ^67^. The same operation was performed for the antisense matrices.

Heatmaps were plotted using plotHeatmap ^67^, setting missing values to white (-- missingDataColor white), and colorMap to Blues (sense) or Reds (antisense). To investigate if the expressed elements contained an intact YY1 binding site, we extracted the relevant sequences using getfasta from bedtools (version 2.30.0; RRID:SCR_006646) ^68^ using GRCh38.p13 as input fasta (-fi) and forcing strandness (-s). We quantified the number of elements with an exact match to the YY1 binding motif (CAAGATGGCCG) ^69^ in the first 100 bp of the element (see github under src/analysis/yy1_present.py).

### PacBio Iso-Seq sample preparation

Total RNA was obtained from tissue samples using miRNA Easy Mini Kit (Qiagen). RNA samples were subsequently put on dry ice and shipped to the National Genomics Infrastructure of Sweden. There, input QC of samples was performed on the Agilent Bioanalyzer instrument, using the Eukaryote Total RNA Nano kit (Agilent) to evaluate RIN and concentration. The sample libraries were prepared as described in “Procedure & Checklist – Iso-Seq™ Express Template Preparation for Sequel® and Sequel II Systems” (PacBio, PN-101763800 Version 02 (October 2019)) using the NEBNext® Single Cell/Low Input cDNA Synthesis & Amplification Module (New England Biolabs, Cat#: E6421S for 24 reactions or E6421L for 96 reactions), the Iso-Seq Express Oligo Kit (PacBio, Cat# PN-101737500), ProNex beads (Promega, Cat#: NG2001 – 10mL, NG2002 – 125mL, NG2003 – 500mL) and the SMRTbell Express Template Prep Kit 2.0 (PacBio, Cat# PN-100938900). 300 ng of total RNA was used for cDNA Synthesis followed by 12 + 3 cycles of cDNA Amplification. In the purification step of amplified cDNA the standard workflow was applied (sample is composed primarily of transcripts centered around 2 kb). After purification the amplified cDNA went into the SMRTbell library construction. Quality control of the SMRTbell libraries was performed with the Qubit dsDNA HS kit (Invitrogen, Cat# Q32851) and the Agilent Bioanalyzer High Sensitivity kit. Primer annealing and polymerase binding was performed using the Sequel II binding kit 2.0 (PacBio, Cat# PN-101789500). Finally, the samples were sequenced on Sequel II and Sequel IIe System using Sequel® II Sequencing Plate 2.0, with an On-Plate Loading Concentration of 110 pM, movie time 24 hours and pre-extension time 2 hours.

Detail protocol can be found at DOI: dx.doi.org/10.17504/protocols.io.yxmvm25j6g3p/v1

For additional information, please contact the National Genomics Infrastructure of Sweden.

#### Iso-Seq mapping to L1HS/PA2 consensus sequence

A L1HS and L1PA2 consensus sequence was used to create a minimap2 (version 2.24; RRID:SCR_018550)^70^ index (minimap2 -d L1consensus.mmi L1consensus.fa) to map FLNC reads (HiFi reads). The density of mapped reads was visualized in the Integrative Genomics Viewer (IgV) (version 2.12.3; RRID:SCR_011793)^71^.

The number of mapped reads in the L1s 5’ UTR was retrieved using samtools view (- c) (version 1.9; RRID:SCR_002105), specifying the first 900 bp of the consensus sequence as the coordinates of interest.

### Isolation of NeuN+ cells

Nuclei were isolated from frozen tissue as described above. Before FACSing, nuclei were incubated with Recombinant Alexa Fluor® 488 Anti-NeuN antibody [EPR12763] Neuronal Marker (Abcam, Cat# ab190195, RRID:AB_2716282) at a concentration of 1:500 for 30 minutes on ice as previously described.^72^ The nuclei were run through the FACS at 4°C with a low flow rate using a 100 mm nozzle and 300.000 nuclei Alexa Fluor – 488 positive nuclei were sorted. The sorted nuclei were pelleted at 1,300 x g for 15 min and resuspended in 1 mL of ice-cold nuclear wash buffer (20 mM HEPES, 150 mM NaCl, 0.5 mM spermidine, 1x cOmplete protease inhibitors, 0.1% BSA) and 10 µL per antibody treatment of ConA-coated magnetic beads (Epicypher) added with gentle vortexing (Pipette tips for transferring nuclei were pre-coated with 1% BSA).

Protocol can be found at DOI: dx.doi.org/10.17504/protocols.io.4r3l27pejg1y/v1.

#### CUT&RUN

We followed the protocol detailed by the Henikoff lab.^41^ Briefly, 100,000 sorted nuclei were washed twice (20 mM HEPES pH 7.5, 150 mM NaCl, 0.5 mM spermidine, 1x Roche cOmplete protease inhibitors) and attached to 10 ConA-coated magnetic beads (Bangs Laboratories) that had been pre-activated in binding buffer (20 mM HEPES pH 7.9, 10 mM KCl, 1 mM CaCl_2_, 1 mM MnCl_2_). Bead-bound cells were resuspended in 50 μL buffer (20 mM HEPES pH 7.5, 0.15 M NaCl, 0.5 mM Spermidine, 1x Roche complete protease inhibitors, 0.02% w/v digitonin, 2 mM EDTA) containing primary antibody (rabbit anti-H3K4me3 Active Motif 39159, RRID:AB_2615077; or goat anti-rabbit IgG, Abcam ab97047, RRID:AB_10681025) at 1:50 dilution and incubated at 4°C overnight with gentle shaking. Beads were washed thoroughly with digitonin buffer (20 mM HEPES pH 7.5, 150 mM NaCl, 0.5 mM Spermidine, 1x Roche cOmplete protease inhibitors, 0.02% digitonin). After the final wash, pA-MNase (a generous gift from Steve Henikoff) was added to the digitonin buffer and incubated with the cells at 4°C for 1 h. Bead-bound cells were washed twice, resuspended in 100 μL digitonin buffer, and chilled to 0-2°C. Genome cleavage was stimulated by the addition of 2 mM CaCl2 at 0°C for 30 min. The reaction was quenched by the addition of 100 μL 2x stop buffer (0.35 M NaCl, 20 mM EDTA, 4 mM EGTA, 0.02% digitonin, 50 ng/μL glycogen, 50 ng/μL RNase A, 10 fg/μL yeast spike-in DNA (a generous gift from Steve Henikoff)) and vortexing. After 10 min of incubation at 37°C to release genomic fragments, cells and beads were pelleted by centrifugation (16,000 g, 5 min, 4°C) and fragments from the supernatant were purified. Illumina sequencing libraries were prepared using the Hyperprep kit (KAPA) (Roche, Cat# 7962347001) with unique dual-indexed adapters (KAPA) (Roche, Cat# 8278555702), pooled and sequenced on a Nextseq500 instrument (Illumina).

Detail protocol can be found at DOI: dx.doi.org/10.17504/protocols.io.j8nlkwb8dl5r/v1.

##### CUT&RUN analysis

Paired-end reads (2x75) were aligned to the human genome (hg38) using bowtie2 (version 2.3.4.2; RRID:SCR_016368) ^73^ (–local –very-sensitive-local –no-mixed –no-discordant –phred33 -I 10 -X 700), converted to bam files with samtools (version 1.4; RRID:SCR_002105) and sorted (samtools version 1.9; RRID:SCR_002105). RPKM normalized bigwig coverage tracks were made with bamCoverage (deepTools (version 2.5.4; RRID:SCR_016366)) ^67^.

Tag directories were created using Homer (version 4.10; RRID:SCR_010881)^74^ makeTagDirectory on default parameters. Peak calling was performed using findPeaks (Homer), using the option histone as style (-style). The rest of the parameters were left on default options. Peaks were then annotated using the script annotatePeaks.pl (Homer; http://homer.ucsd.edu/homer/ngs/annotation.html) and intersected (BEDtools, version 2.30.0; RRID:SCR_006646) to bed files containing coordinates of >6kbp L1HS, L1PA2, L1PA3 or L1PA4. Matrices for heatmaps were created (computeMatrix, deeptools, version 2.5.4; RRID:SCR_016366) using the peaks with an overlap on these elements (only peaks which were called in all samples of a dataset) and visualized using plotHeatmap (deeptools).

## Data and code availability

There are no restrictions on data availability. The RNA and DNA sequencing data presented in this study have been deposited at GEOs:

- GSE225081: Bulk, short and long read, RNAseq of adult samples. We also included Cell Ranger’s matrices that were used for this paper from the 5’ 10x snRNAseq (raw data has been previously published at GSE211870).
- GSE224747: 3’ single nuclei RNAseq, CUT&RUN and bulk RNAseq of fetal samples.
- GSE224659: CRISPRi in fbNPCs and organoids.

Any additional information required to reanalyze the data reported in this paper is available from the lead contact upon request.

This paper analyzes existing, publicly available data. These accession numbers for the datasets are:

- GSE209552: 3’ single nuclei RNAseq of adult samples^75^.
- GSE211870: 5’ single nuclei RNAseq of adult samples^76^.
- GSE211871: Adult Neun+ CUT&RUN^76^.
- GSE182224: Chimpanzee and human fbNPCs^46^.
- original code has been deposited at GitHub and is publicly available at: https://github.com/raquelgarza/truster.git(doi:10.5281/zenodo.7589548) and https://github.com/raquelgarza/L1_transcriptional_complexity_Garza2023.git (doi:10.5281/zenodo.7696265)

## Author contributions

All of the authors took part in designing the study and interpreting the data. R.G., D.A. and J.J. conceived and designed the study. D.A., A.A., P.G., O.K., V.H., P.A.J., M.V., J.M., M.J., E.M. and C.H.D. performed the experimental research. R.G., D.A., N.P., P.H., Y.S. and C.H.D. performed the bioinformatic analyses. A.Q., A.K., N.M., E.E., E.E.E., M.H., R.A.B. and Z.K. contributed material, reagents, and expertise. R.G., D.A. and J.J. wrote the manuscript, and all authors reviewed the final version.

## Acknowledgments

We would like to thank Jonas Frisén, Didier Trono, Fred H. Gage, Wieland Huttner, Svante Pääbo, Niklas Marklund and Steve Henikoff, for their comments and support. We also thank U. Jarl, M. Persson Vejgården and A. Hammarberg for their technical assistance. We are grateful to all members of the Jakobsson lab. The work was supported by grants from the Aligning Science Across Parkinson’s (ASAP-000520 to J.J.) through the Michael J. Fox Foundation for Parkinson’s Research, the Swedish Research Council (2018-02694, to J.J. ; 2020-01660 to Z.K and 2021-03494 to C.H.D.), the Swedish Brain Foundation (FO2019-0098, to J.J., FO2022-0079 to Z.K.), Cancerfonden (190326, to J.J.), Barncancerfonden (PR2017-0053, to J.J.), NIHR Cambridge Biomedical Research Centre (NIHR203312, to R.A.B) the Swedish Society for Medical Research (S19-0100, to C.H.D.), National Institutes of Health (HG002385 to E.E.E.) and the Swedish Government Initiative for Strategic Research Areas (MultiPark & StemTherapy). E.E.E. is an investigator of the Howard Hughes Medical Institute. For the purpose of open access, the author has applied a CC BY public copyright licence to all Author Accepted Manuscripts arising from this submission.

---

**Supplemental Figure 1.**
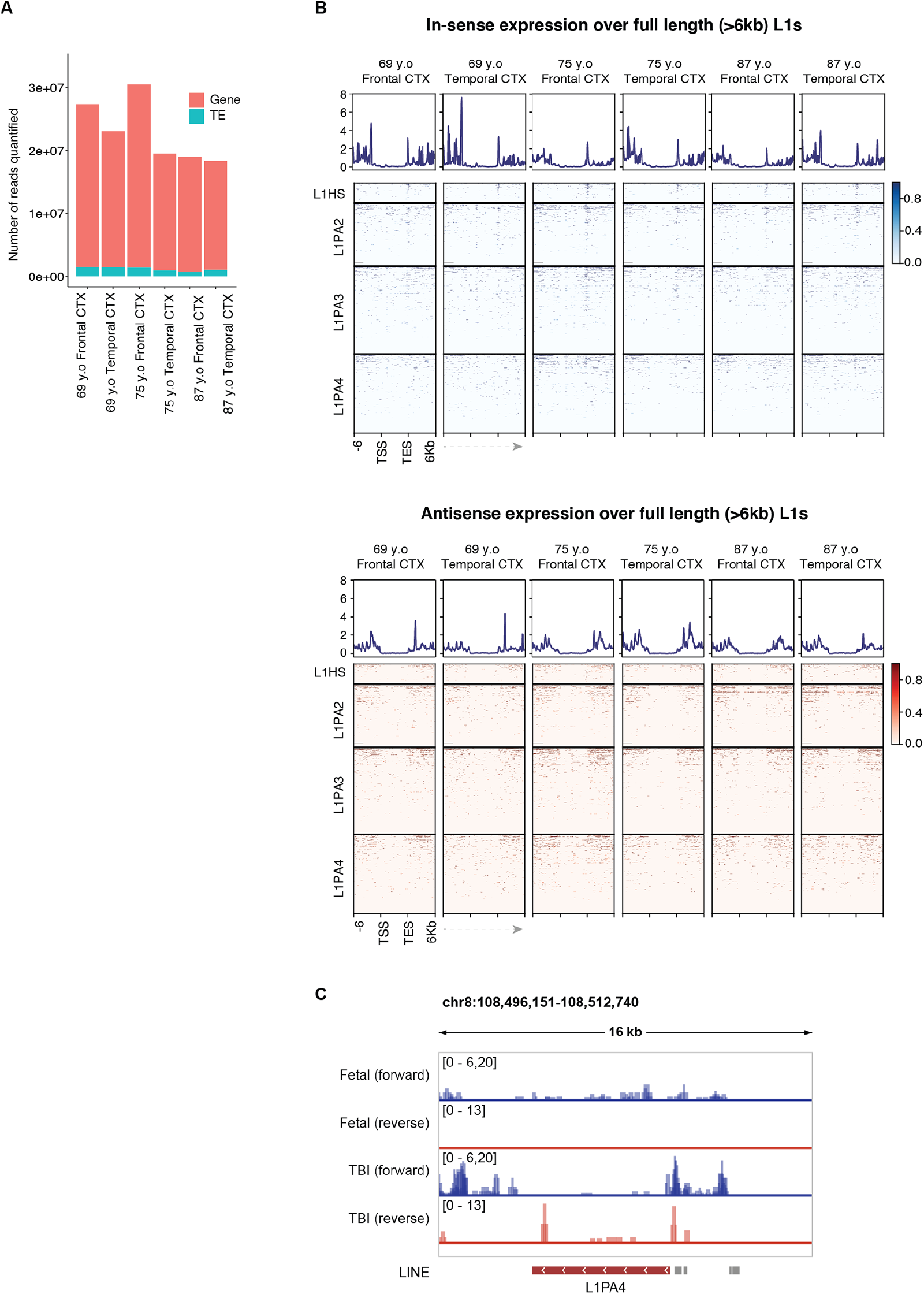
A) Number of reads quantified as genes or TEs per sample, as quantified by TEcounts. B) Expression (RPKM) over full length (>6kbp) L1HS, L1PA2, L1PA3 and L1PA4, plus 6kbp flanking regions. Blue heatmaps showing the signal per sample in sense of the annotated element. Red heatmaps showing signal in antisense. C) Genome browser tracks showing an adult-specific expression of a >6kbp L1PA4 with antisense transcription initiated in its promoter.

**Supplemental Figure 2.**
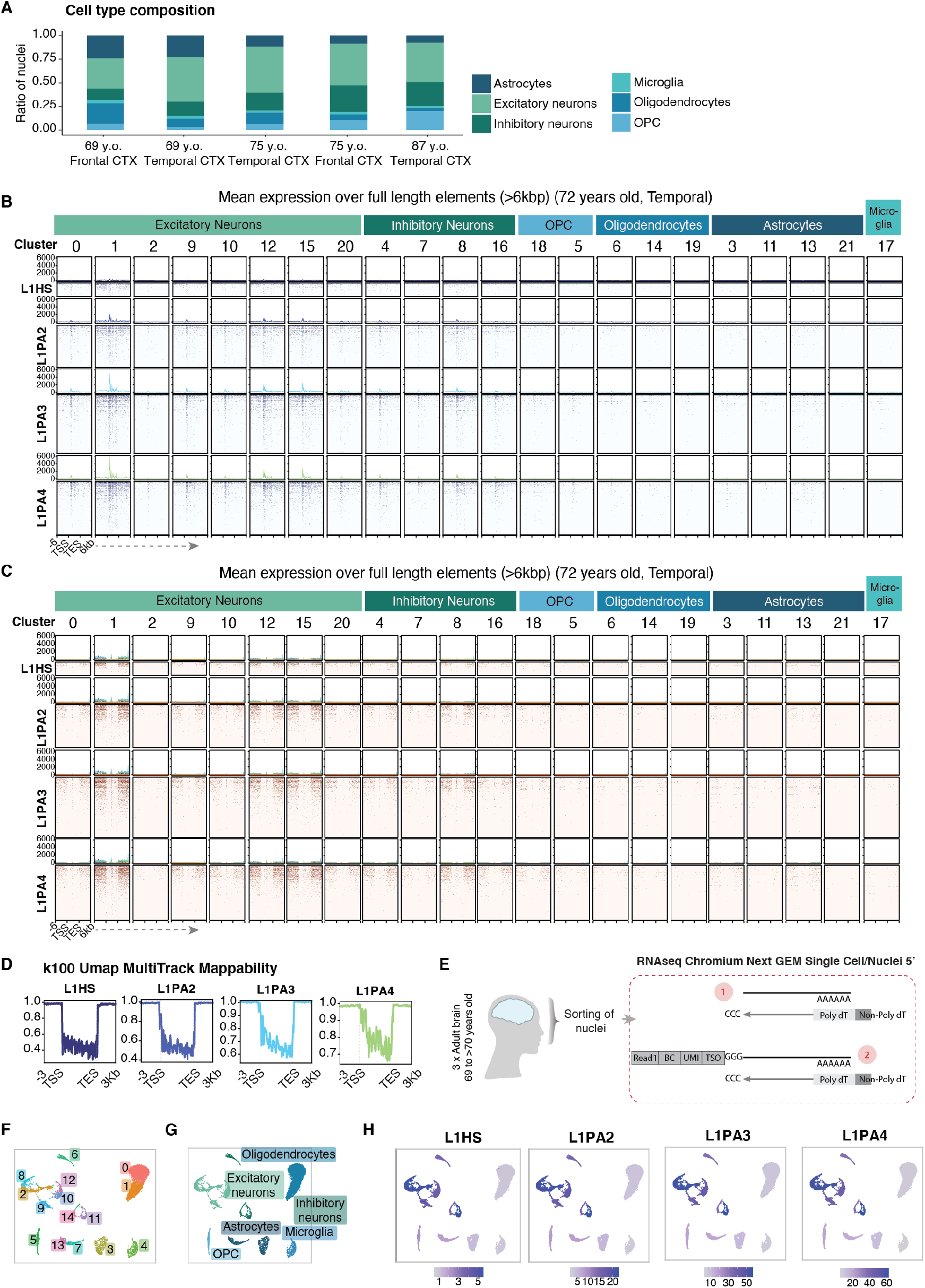
A) Cell type composition in the single nuclei RNAseq of adult samples. B) Expression (RPKM) over full length (>6kbp) L1HS, L1PA2, L1PA3 and L1PA4, plus 6kbp flanking regions in each cluster for one of the adult samples. Blue heatmaps showing the signal per cluster in sense of the annotated element. Red heatmaps showing signal in antisense. Top annotation indicates the cell type of the cluster in question. D) Single-read mappability score for full-length (>6kbp) young L1 subfamilies (read length of 100) as reported for hg38 by Karimzadeh, et al. 2018 (tracks available at UCSC table browser)). E) Schematic of 5’ enrichment Chromium Next GEM library. F) single nuclei RNAseq UMAP colored by cluster. G) UMAP colored by characterized cell types. H) Pseudo-bulk cluster expression of young L1 subfamilies on UMAP.

**Supplemental Figure 3.**
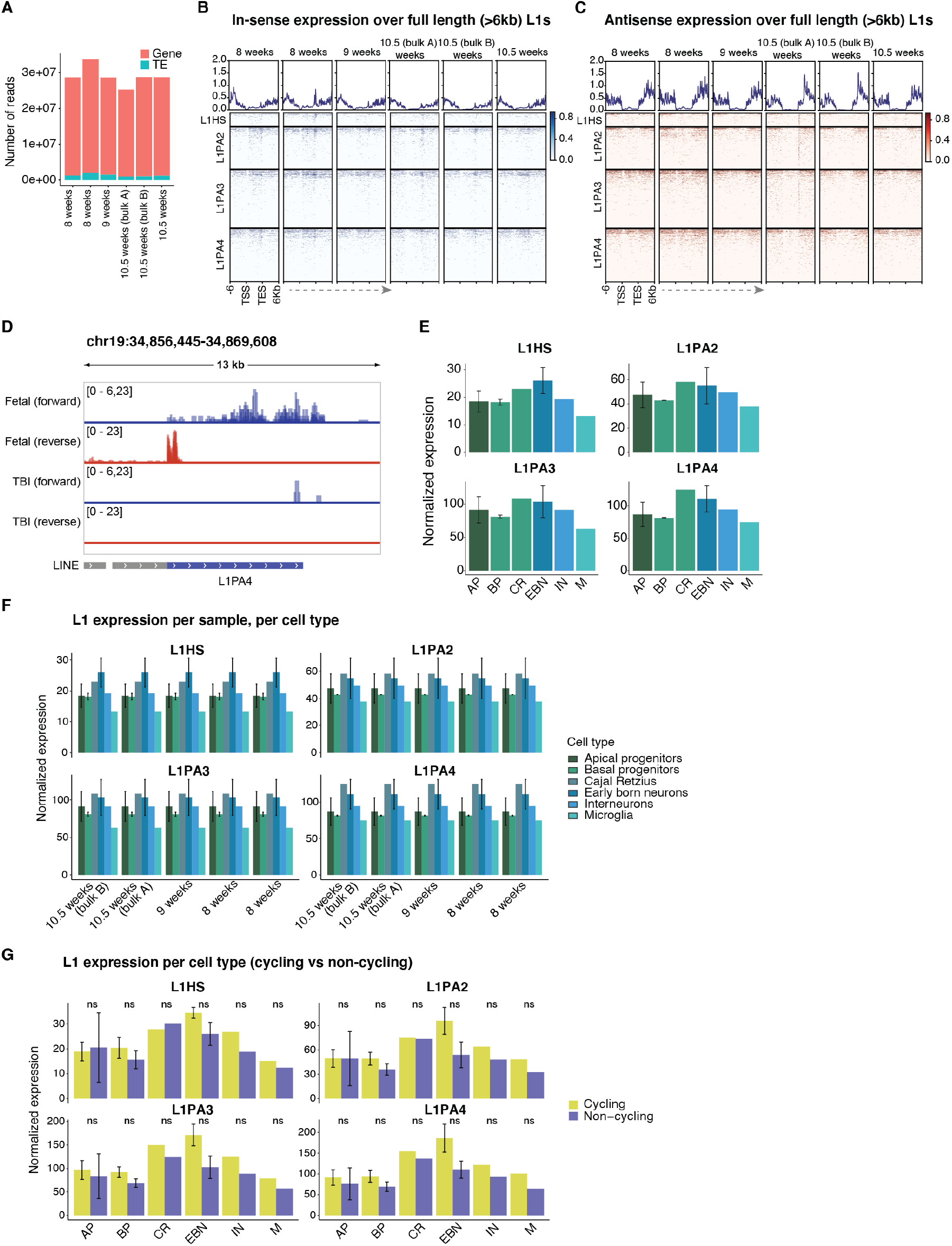
A) Number of reads quantified as genes or TEs per sample, as quantified by TEcounts. B) Expression (RPKM) over full length (>6kbp) L1HS, L1PA2, L1PA3 and L1PA4, plus 6kbp flanking regions. Blue heatmaps showing the signal per sample in sense of the annotated element. C) Red heatmaps showing signal in antisense D) Genome browser tracks showing an fetal-specific expression of a >6kbp L1PA4 with antisense transcription initiated in its promoter. E) Comparison of the pseudo-bulk cluster expression of young L1 subfamilies among the different cell types (AP = apical progenitors; BP = basal progenitors; CR = Cajal Retzius; EBN = early-born neurons; IN = interneurons; M = microglia). F) Cluster expression of young L1 subfamilies (quantified per sample), grouped per cell type. G) L1 expression of cycling vs non-cycling cells from each cluster, grouped per cell type (paired Wilcoxon test).

**Supplemental Figure 4.**
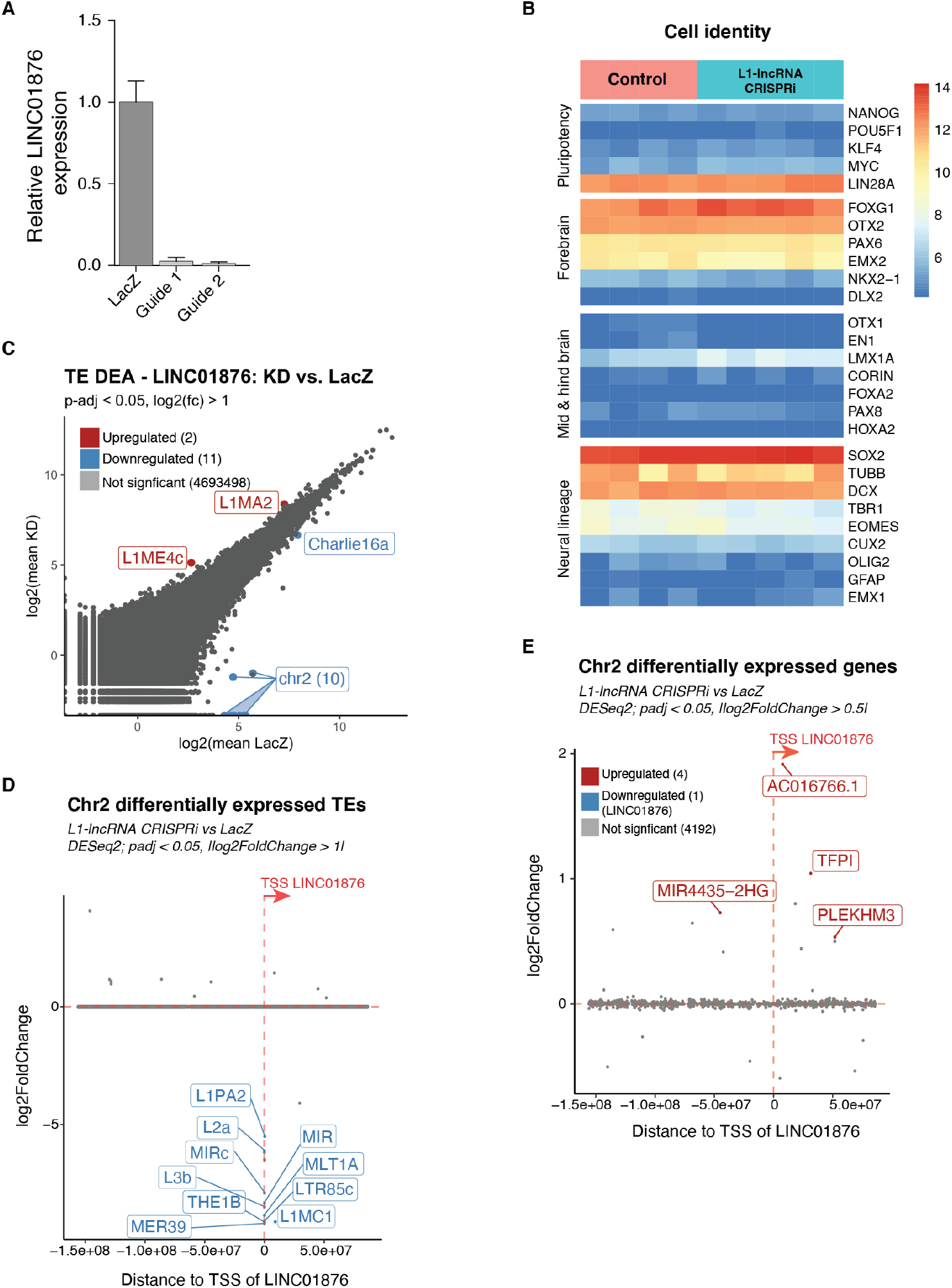
A) qPCR of L1-lncRNA in control (LacZ) and L1-lncRNA CRISPRi guide 1 and 2. B) Selected gene markers to represent cell identity in controls and L1-lncRNA fbNPCs. C) Mean plot showing results of differential expression analysis of TEs (DESeq2). Significantly upregulated elements (padj < 0.05; log2FoldChange >1) highlighted in red; significantly downregulated elements (padj < 0.05; log2FoldChange < -1) highlighted in blue. Labels showing TE subfamily or if the TE is located in chr2. Manhattan plot of chr2 showing log2FoldChange of TEs. Differentially expressed TEs nearby L1-lncRNA (start site at x = 0) are highlighted in red or blue (up and downregulated, respectively) (DESeq2, |log2FoldChange > 1|; padj < 0.05). D) Manhattan plot of chr2 showing log2FoldChange of genes. Differentially expressed genes nearby L1-lncRNA (start site at x = 0) are highlighted in red and blue (up and downregulated, respectively) (DESeq2, log2FoldChange > 0.5; padj < 0.05).

**Supplemental Figure 5.**
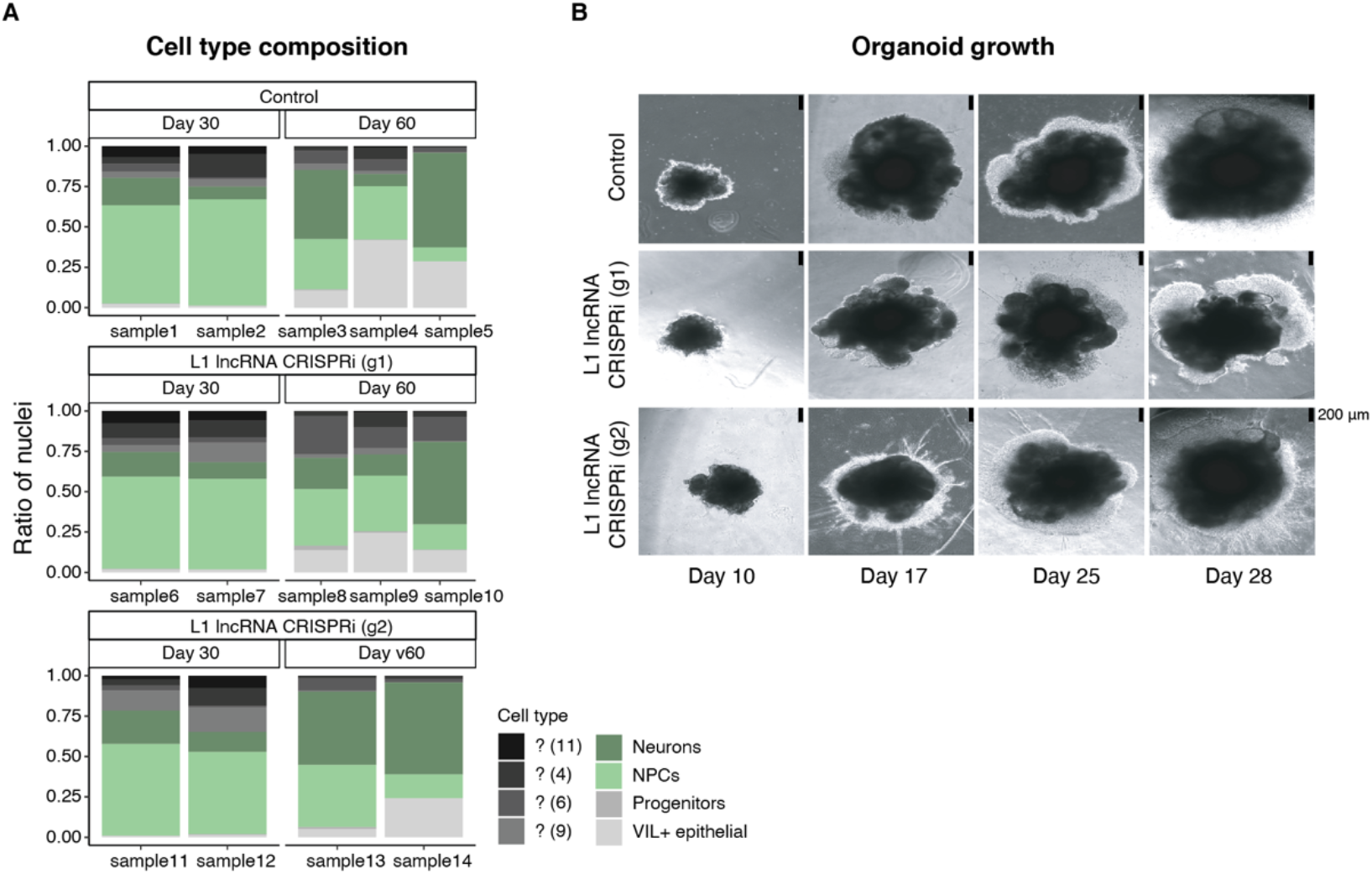
A) Cell type composition in control (LacZ) and L1-lncRNA CRISPRi (g1 and g2) cerebral organoids. Neural-like cell types colored in green. B) Brightfield imaging showing organoids sizes at day 10, 17, 25 and 28 (scale 200 μm, black bar) in control (LacZ) and L1-lncRNA CRISPRi (g1 and g2).

**Supplemental Figure 6.**
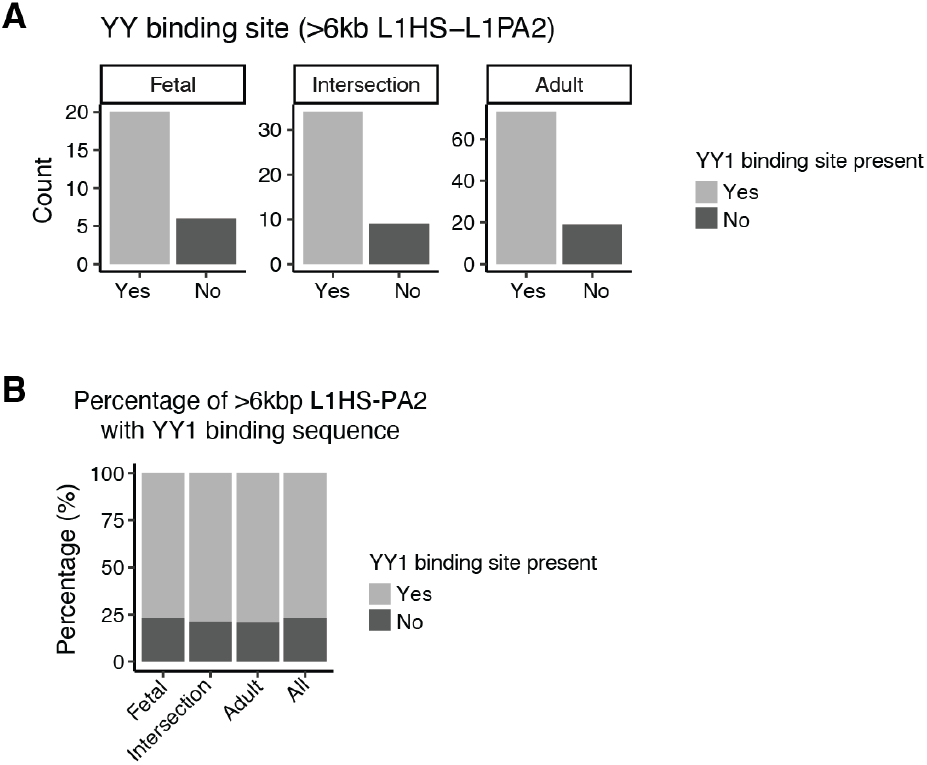
A) Number of expressed L1HS-L1PA2 (>6kbp) with YY1 binding sequence present (see methods) in fetal, adult, or those expressed in both datasets (intersection). B) Percentage of >6kbp L1HS-L1PA2 with YY1 binding sequence among those expressed in fetal samples, adult samples, both datasets (intersection), and all annotated in hg38 (see methods).

